# Serum albumin restrains obesity by coordinating brown adipose thermogenic capacity with β3-adrenergic responsiveness

**DOI:** 10.64898/2026.11.20.767998

**Authors:** Zongrui Luo, Yi Jiang, Hongyi Liu, Xuan Dong, Enkai Li, Jingyi Zhu, Yan Fu, Yongzhang Luo

## Abstract

Obesity reflects chronic imbalance between energy storage and expenditure, yet how abundant circulating carrier proteins contribute remains poorly understood. Using constitutive and inducible albumin-deficient mice, we show that reduced circulating albumin promotes diet-induced adiposity and hepatic lipid accumulation while impairing energy expenditure, cold tolerance and stimulus-induced lipid mobilization. Recombinant mouse serum albumin (rMSA) restores thermogenic capacity and alleviates obesity, with anti-adiposity effects partly dependent on UCP1. Mechanistically, rMSA promotes brown-adipocyte thermogenic maturation through Cav1-associated Src–PI3K/AKT–mTORC1 signaling. Independently, the fatty-acid-to-albumin balance tunes β3-adrenergic responsiveness: high fatty-acid loading shifts Cav1-associated complexes from β3-AR toward CD36, whereas CD36 depletion restores β3-AR association, PKA/HSL signaling and glycerol release. Glycation or oxidation impairs albumin fatty-acid binding and is associated with reduced metabolic efficacy. Thus, albumin abundance, fatty-acid loading and biochemical integrity jointly shape brown-adipose thermogenesis and obesity susceptibility.

## INTRODUCTION

Obesity, defined as having a body mass index (BMI) of 30 kg/m^2^ or higher, has become increasingly prevalent worldwide in the past four decades due to changes in diet and increasing sedentary behavior^1,2^. The World Health Organization (WHO) describes obesity as a major public health issue since it increases the risk of multiple diseases, including type 2 diabetes mellitus (T2DM), cardiovascular disease, hypertension, nonalcoholic fatty liver disease, and certain types of cancer^3,4^. Bariatric surgery is currently the most effective method of achieving weight loss^5^. However, surgical interventions are not capable of addressing the global demand for weight loss and may be harmful to the body^6,7^. Currently, several pharmacological agents have been approved by the US FDA for chronic weight management, including orlistat and incretin-based therapies such as the glucagon-like peptide-1 receptor (GLP-1R) agonists liraglutide and semaglutide^6,8^. However, these therapies are frequently associated with gastrointestinal adverse effects, including nausea, vomiting, and diarrhea, and discontinuation of GLP-1R agonist treatment can result in substantial weight regain^9,10^. Therefore, despite recent advances in anti-obesity pharmacotherapy, it remains critical to unravel the mechanisms underlying obesity and develop effective therapeutics.

Obesity, characterized by excessive fat accumulation, is a complex chronic disease that arises from an imbalance between energy intake and expenditure and is influenced by genetic, physiological, and environmental factors^11,12^. Overexpansion of white adipose tissue (WAT) caused by excess energy storage leads to metabolic disorders associated with obesity^13^. There is another type of adipose tissue, brown adipose tissue (BAT), which serves a distinct function from WAT^14^. Brown adipocytes are smaller and store triglycerides in multilocular droplets compared to unilocular ones found in white fat ^15^. In addition, brown adipocytes contain many mitochondria and express high levels of uncoupling protein 1 (UCP1), which mediates thermogenesis by separating substrate oxidation from ATP synthesis into heat production^16,17^. Physiologically, brown adipose thermogenesis is triggered by cold stimulation or dietary intake, and it involves the regulation of the PKA-p38-Ucp1 signaling axis^18^. During cold exposure, sympathetic adrenergic signaling stimulates adipose lipolysis, in rodents predominantly through β3-adrenergic receptors (β3-ARs), thereby mobilizing fatty acids that contribute to the fuel supply for adaptive thermogenesis. The insulin signaling pathway is critical for brown adipocyte development, and mTORC1 activation has been reported to profoundly regulate the thermogenic function of brown adipose tissue^19,20^. Previous studies suggested that UCP1-mediated thermogenesis may be beneficial for treating obesity in both human and mouse models^21,22^. Therefore, discovering promising gene targets and approaches toward safe and effective brown fat activation is urgently required.

Albumin, which accounts for about 60% of total protein in serum, plays a variety of biological functions, including maintaining plasma oncotic pressure and serving as a major carrier of non-esterified fatty acids in the circulation^23,24^. Low serum albumin levels have been observed in obese individuals^25–27^. In fact, the biological functions of serum albumin can be indirectly elucidated through limited studies on congenital analbuminemia, a rare genetic disorder in humans ^28^. In the limited available case reports, analbuminemic individuals consistently present with notable hypercholesterolemia accompanied by a distinct clinical presentation of enlarged lower extremities^29,30^. Our group previously demonstrated that long-term administration of young, undamaged, and ultrapure recombinant mouse serum albumin (rMSA) can extend the healthy lifespan of mice without side effects^31^. Moreover, rMSA has also been shown to be a potential treatment for T2DM^32^. Given the important metabolic functions of albumin, we sought to determine whether circulating albumin regulates the development of obesity and to define the mechanisms underlying its metabolic actions.

Here, using constitutive and inducible albumin-deficient mouse models, we show that circulating albumin restrains diet-induced adiposity and supports adaptive thermogenesis and stimulus-induced lipid mobilization. Mechanistically, rMSA promotes brown-adipocyte thermogenic maturation through Cav1-associated Src–PI3K/AKT–mTORC1 signaling, whereas the fatty-acid-to-albumin balance tunes β3-adrenergic responsiveness through Cav1/CD36-associated membrane remodeling. Albumin glycation and oxidation impair fatty-acid binding and reduce metabolic efficacy. Together, these findings identify albumin abundance, fatty-acid loading state and biochemical integrity as complementary determinants of brown-adipose thermogenic function and obesity susceptibility.

## RESULTS

### Albumin deficiency exacerbates HFD-induced obesity

To determine whether a reduction in circulating albumin is sufficient to accelerate diet-induced obesity while minimizing potential developmental confounding, we generated an inducible albumin-knockout (iAKO) model by administering Cre-expressing AAV to *Alb^fl/fl^* mice, with AAV-Cre-treated C57BL/6J mice serving as WT controls (Fig. 1A). AAV-Cre administration markedly reduced circulating albumin (Fig. S1A). When challenged with a high-fat diet (HFD), iAKO mice gained significantly more body weight than WT controls (Fig. 1B). EchoMRI analysis revealed greater fat mass and fat percentage in iAKO mice (Fig. 1C). Absolute lean mass remained unchanged, whereas lean mass as a percentage of body weight was reduced in iAKO mice, consistent with the disproportionate increase in adiposity (Fig. 1D).

**Figure 1.**
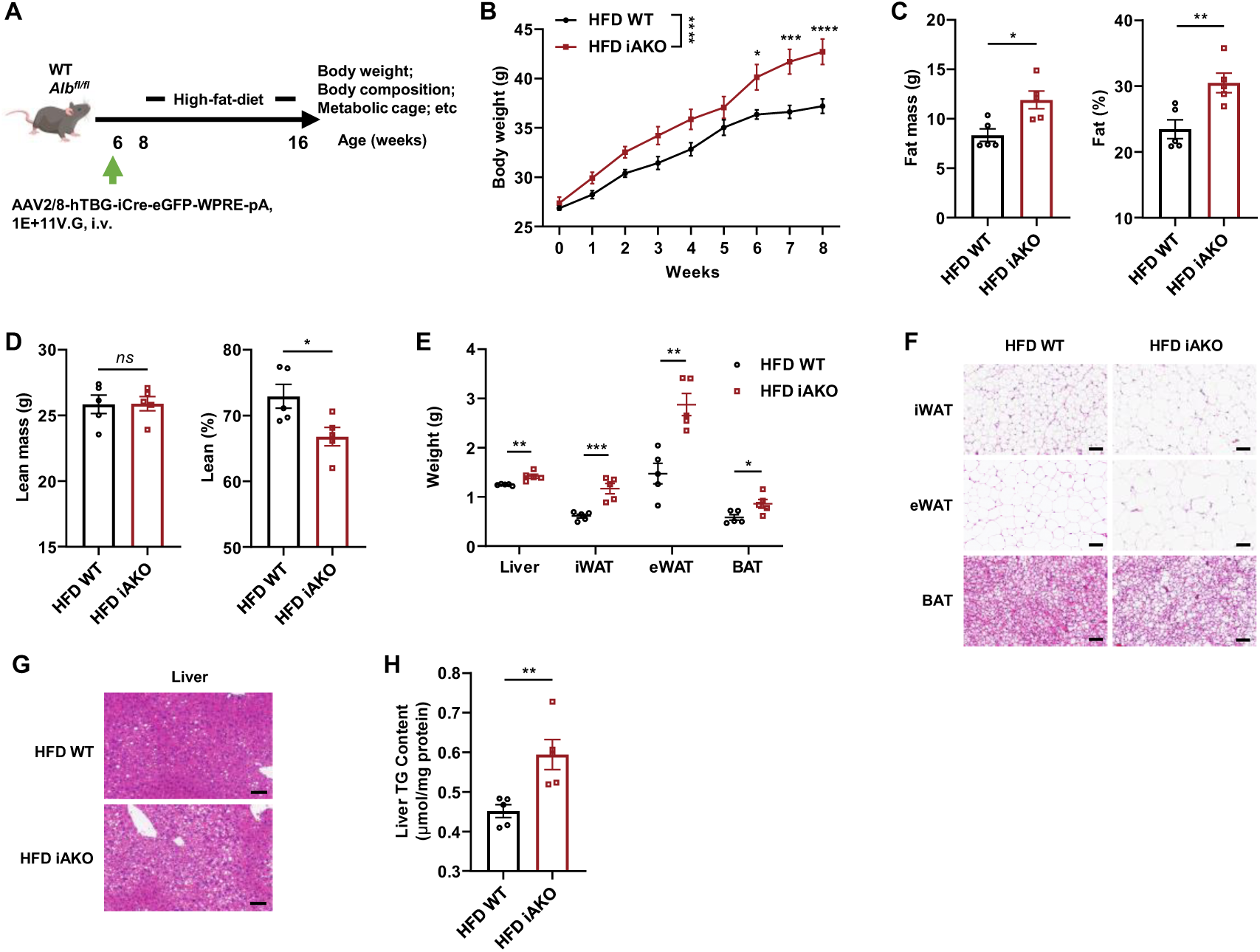
Albumin deficiency exacerbates obesity progression in mice. (A) Schematic illustration of the animal experimental procedure. (B) Body weights of WT and inducible Albumin knockout (iAKO) mice on HFD. n = 5 mice per group. (C) Fat mass (left) and fat percentage (right) of WT and iAKO mice on HFD. n = 5 mice per group. (D) Lean mass (left) and lean percentage (right) of WT and iAKO mice on HFD. n = 5 mice per group. (E) Weights of liver and adipose tissue. n = 5 mice per group. (F) Representative H&E staining of dissected adipose tissues. Scale bar, 100 μm. (G) Representative H&E staining of dissected liver tissues. Scale bar, 100 μm. (H) Quantification of liver TG content. n = 5 mice per group. Data are presented as mean ± s.e.m; \**P* < 0.05, \*\**P* < 0.01, \*\*\**P* < 0.001, \*\*\*\**P* < 0.0001 compared to control group. *ns*, not significant; two-tailed Student’s t-tests (C-E, and H), two-way ANOVA followed by Sidak’s multiple comparisons (B).

Consistent with the body-composition phenotype, iAKO mice had increased liver and adipose-tissue weights (Fig. 1E). Histological analysis revealed enlarged white adipocytes, whitening of brown adipose tissue (BAT), and greater hepatic lipid accumulation (Fig. 1F, G). Hepatic triglyceride (TG) content was also significantly elevated in iAKO mice (Fig. 1H). A constitutive albumin-knockout model independently recapitulated this phenotype, including accelerated HFD-induced weight gain, increased fat mass, reduced relative lean mass, expansion of adipose depots, adipocyte hypertrophy, BAT whitening, and hepatic steatosis (Fig. S1C–K). Together, these loss-of-function models demonstrate that reduced albumin availability increases susceptibility to HFD-induced adiposity and ectopic lipid accumulation.

### Restoration of circulating albumin ameliorates the obesity phenotype of iAKO mice

We next asked whether restoring circulating albumin could ameliorate the metabolic consequences of albumin depletion. HFD-fed iAKO mice were treated with recombinant mouse serum albumin (rMSA), which restored circulating albumin toward the physiological range (Fig. 2A and Fig. S1B). Albumin replacement substantially attenuated HFD-induced weight gain in iAKO mice (Fig. 2B). rMSA also reduced fat mass and fat percentage, while absolute lean mass remained largely unchanged and relative lean mass increased (Fig. 2C, D).

**Figure 2.**
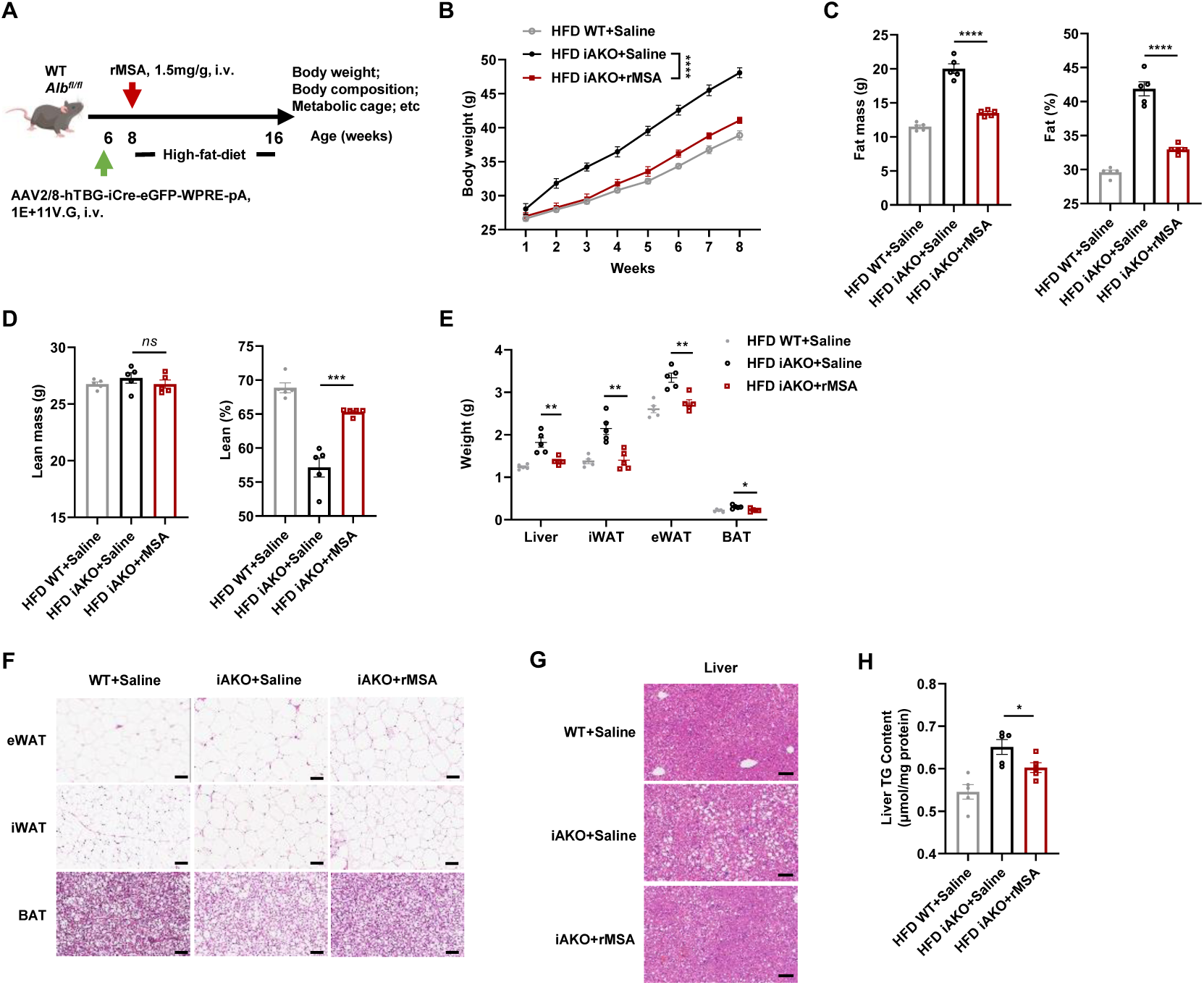
Changes in serum albumin levels regulate obesity progression in mice. (A) Schematic illustration of the animal experimental procedure. (B) Body weights of WT treated with saline and iAKO mice treated with saline or rMSA (1.5 mg/g body weight) on HFD. n = 5 mice per group. (C) Fat mass (left) and fat percentage (right) of WT treated with saline and iAKO mice treated with saline or rMSA on HFD. n = 5 mice per group. (D) Lean mass (left) and lean percentage (right) of WT treated with saline and iAKO mice treated with saline or rMSA on HFD. n = 5 mice per group. (E) Weights of liver and adipose tissue. n = 5 mice per group. (F) Representative H&E staining of dissected adipose tissues. Scale bar, 100 μm. (G) Representative H&E staining of dissected liver tissues. Scale bar, 100 μm. (H) Quantification of liver TG content. n = 5 mice per group. Data are presented as mean ± s.e.m; \**P* < 0.05, \*\**P* < 0.01, \*\*\**P* < 0.001, \*\*\*\**P* < 0.0001 compared to control group. *ns*, not significant; one-way ANOVA (C-E, and H), two-way ANOVA followed by Sidak’s multiple comparisons (B).

These changes in body composition were accompanied by lower liver and adipose-tissue weights (Fig. 2E), smaller adipocytes and reduced BAT whitening (Fig. 2F), and alleviation of hepatic steatosis (Fig. 2G). Consistently, hepatic TG content was reduced by rMSA treatment (Fig. 2H). Thus, restoring circulating albumin substantially ameliorates the adiposity and hepatic lipid accumulation caused by albumin depletion.

### Albumin deficiency alters systemic glucose homeostasis

Because obesity is commonly accompanied by impaired glucose homeostasis, we examined whether the increased adiposity associated with albumin deficiency was accompanied by a corresponding deterioration in glucose tolerance. Unexpectedly, constitutive AKO mice exhibited lower glucose excursions during glucose tolerance testing than WT controls, whereas insulin tolerance was only modestly affected (Fig. S2A, B). A similar phenotype was observed in iAKO mice, which showed reduced glucose excursions during GTT and a lower ITT AUC than WT mice (Fig. S2C, D). Thus, despite their greater adiposity, albumin-deficient mice did not exhibit the glucose-intolerant phenotype typically associated with obesity.

Notably, fasting glucose measured before glucose administration during GTT was already lower in albumin-deficient mice, raising the possibility that the reduced glycemic excursions partly reflected a lower baseline glucose level. Consistently, AKO mice exhibited lower blood glucose concentrations after both short and prolonged fasting (Fig. S2G). We therefore examined endogenous glucose production. iAKO mice showed a blunted glycemic response to pyruvate, whereas rMSA administration restored this response toward that of WT mice (Fig. S2H). However, although rMSA replacement restored circulating albumin in HFD-fed iAKO mice toward physiological levels, it did not significantly alter GTT or ITT AUC (Fig. S2E, F). Moreover, the difference in baseline-adjusted GTT between WT and iAKO mice persisted (Fig. S2I). Collectively, these findings suggest that the altered glucose-handling phenotype associated with albumin deficiency is not explained solely by lower fasting glucose and is not fully normalized by restoration of circulating albumin alone, revealing an unexpected dissociation between adiposity and glucose handling in albumin-deficient mice.

### Serum albumin regulates energy homeostasis through modulation of thermogenesis

BAT plays a critical role in energy metabolism regulation due to its unique thermogenic capacity^33^. Having established that albumin deficiency promotes adiposity and that this phenotype can be substantially ameliorated by restoring circulating albumin, we next sought to define the physiological basis for these changes in energy balance. Because excessive fat accumulation can result from increased energy intake and/or reduced energy expenditure, we examined whole-body energy metabolism in HFD-fed iAKO mice. Whole-body oxygen consumption (VO2) was reduced in HFD-fed iAKO mice relative to WT controls (Fig. 3A), whereas daily food intake and locomotor activity were unchanged (Fig. 3B). Conversely, rMSA treatment increased VO2 in iAKO mice without altering food intake or locomotor activity (Fig. 3C, D), arguing against differences in food intake or locomotor activity as major explanations for the observed metabolic phenotype.

**Figure 3.**
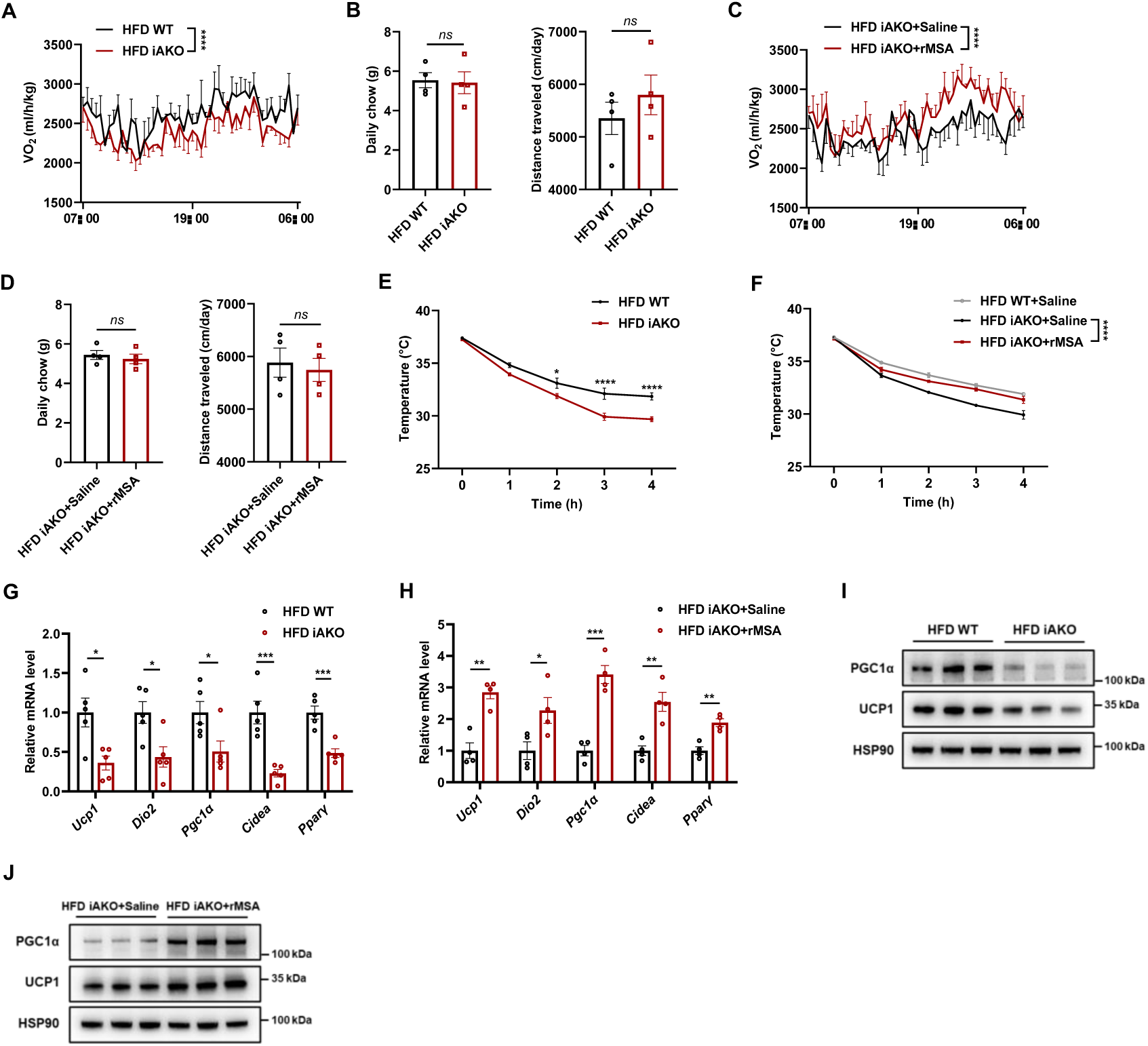
Serum albumin regulates energy homeostasis and thermogenesis. (A) Whole-body VO_2_ normalized by body weight of WT and iAKO mice. n = 4 mice per group. (B) Daily chow and distance traveled of WT and iAKO mice. n = 4 mice per group. (C) Whole-body VO_2_ normalized by body weight of iAKO mice treated with saline or rMSA. n = 4 mice per group. (D) Daily chow and distance traveled of iAKO mice treated with saline or rMSA. n = 4 mice per group. (E) Body temperature of WT and iAKO mice exposed to cold. n = 5 mice per group. (F) Body temperature of WT mice treated with saline and iAKO mice treated with saline or rMSA under cold exposure. n = 5 mice per group. (G) Abundance of thermogenesis-related mRNAs in BAT. n = 5 mice per group. (H) Abundance of thermogenesis-related mRNAs in BAT. n = 4 mice per group. (I) Immunoblots of thermogenesis-related proteins in BAT. n = 3 biological replicates per group. (J) Immunoblots of thermogenesis-related proteins in BAT. n = 3 biological replicates per group. Data are presented as mean ± s.e.m; \**P* < 0.05, \*\**P* < 0.01, \*\*\**P* < 0.001, \*\*\*\**P* < 0.0001 compared to control group. *ns*, not significant; two-tailed Student’s t-tests (B, D, G, and H), two-way ANOVA followed by Sidak’s multiple comparisons (A, C, E, and F).

Given the reduction in whole-body oxygen consumption, we next asked whether adaptive thermogenesis was impaired. During acute cold exposure, iAKO mice exhibited a more rapid decline in core body temperature than WT mice (Fig. 3E), whereas rMSA replacement improved cold tolerance in iAKO mice (Fig. 3F). At the molecular level, BAT from iAKO mice showed reduced expression of thermogenic genes, including *Ucp1*, *Dio2*, *Pgc1α*, *Cidea*, and *Pparγ* (Fig. 3G), while rMSA treatment increased expression of these genes in iAKO BAT (Fig. 3H). Corresponding changes were also evident at the protein level: The abundance of PGC1α and UCP1 was reduced in iAKO BAT and increased after rMSA treatment (Fig. 3I, J).

To further investigate the molecular basis underlying impaired adaptive thermogenesis in albumin-deficient mice, we performed RNA sequencing of BAT from HFD-fed WT and AKO mice. Principal component analysis showed clear separation of the BAT transcriptomic profiles between WT and AKO mice (Fig. S3A). KEGG pathway over-representation analysis of genes significantly downregulated in AKO BAT revealed enrichment of thermogenesis, oxidative phosphorylation, and multiple carbon- and energy-metabolism pathways, including carbon metabolism, propanoate metabolism, pyruvate metabolism, and glycolysis/gluconeogenesis (Fig. S3B). Consistently, preranked GSEA showed significant enrichment of the KEGG thermogenesis and oxidative phosphorylation gene sets toward the WT-associated end of the ranked gene list (Fig. S3C). These transcriptional alterations were accompanied by reduced whole-body oxygen consumption and energy expenditure without significant changes in food intake or locomotor activity (Fig. S3D–F), impaired cold tolerance (Fig. S3G, H), reduced thermogenic gene expression in adipose tissues, decreased PGC-1α and UCP1 protein abundance, and diminished UCP1 staining in BAT (Fig. S3I–L). Together, these findings indicate that albumin deficiency suppresses thermogenic, mitochondrial oxidative, and energy-metabolic programs in adipose tissue.

### rMSA administration limits obesity progression in DIO mice

Having established that albumin depletion aggravates obesity and that albumin restoration ameliorates the phenotype in iAKO mice, we next asked whether exogenous albumin administration could limit obesity progression in mice with established obesity. Consistent with the potential relevance of albumin to obesity, circulating albumin levels were reduced in obese mice (Fig. S4A–C). DIO mice were therefore treated with rMSA beginning after prolonged HFD feeding (Fig. 4A). The young and undamaged rMSA used in this study was the same as that in our previous study ^31,32^. rMSA significantly slowed subsequent body-weight gain (Fig. 4B, C), reduced fat mass and fat percentage (Fig. 4D), and increased the relative contribution of lean mass without changing absolute lean mass (Fig. 4E). Liver, iWAT, eWAT, and BAT weights were lower in rMSA-treated DIO mice (Fig. 4F), and histological analysis showed reduced adipocyte hypertrophy, reduced BAT lipid accumulation, and less hepatic steatosis (Fig. 4G).

**Figure 4.**
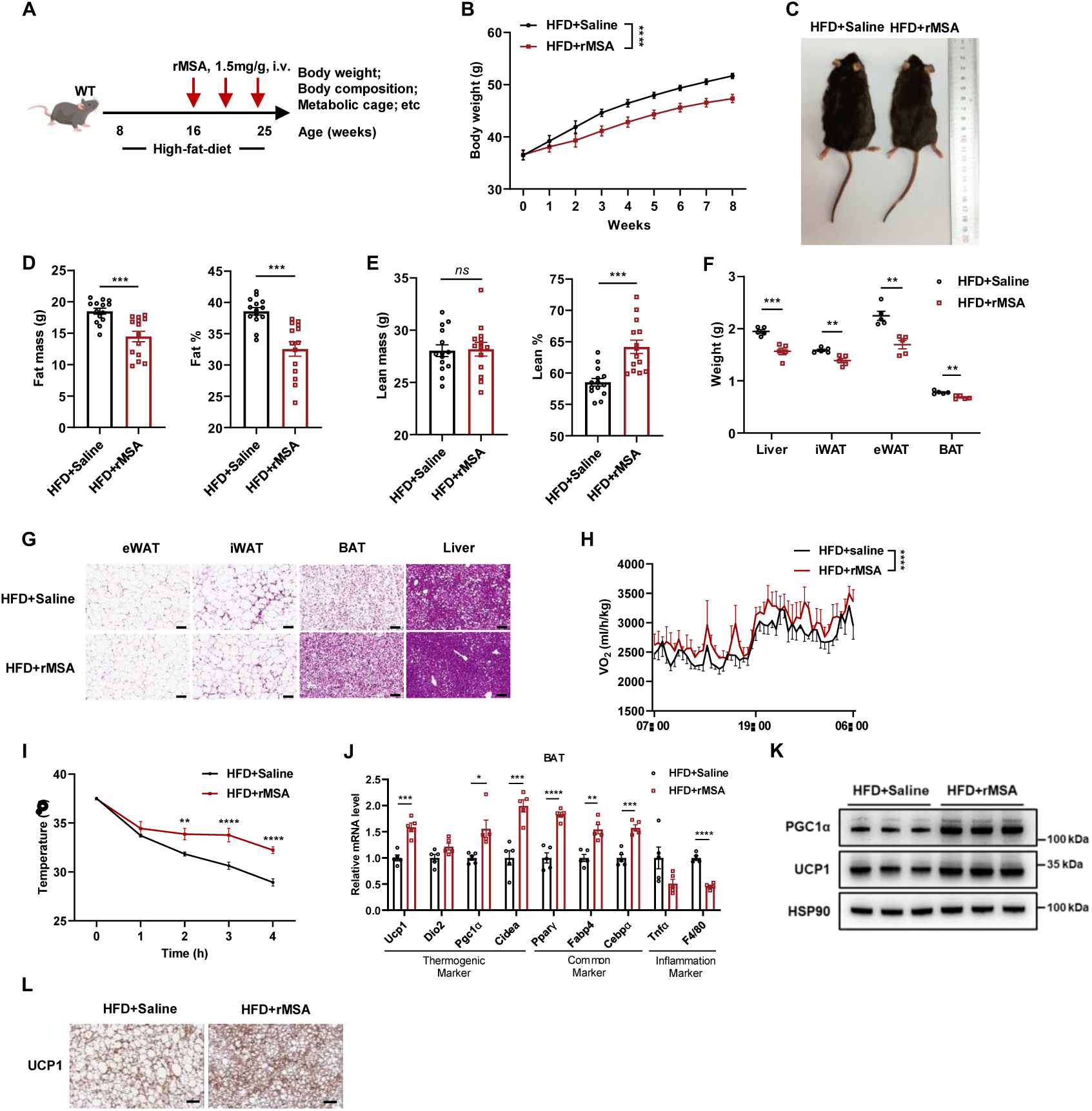
rMSA administration protects against obesity. (A) Schematic illustration of animal experimental procedure. (B) Body weights of DIO mice treated with saline or rMSA. n = 14 mice per group. (C) Representative photographs of DIO mice treated with saline or rMSA. (D) Fat mass (left) and fat percentage (right) of DIO mice treated with saline or rMSA. n = 14 mice per group. (E) Lean mass (left) and lean percentage (right) of DIO mice treated with saline or rMSA. n = 14 mice per group. (F) Weights of liver and fat tissue. n = 5 mice per group. (G) Representative H&E staining of dissected tissues in DIO mice treated with saline or rMSA. Scale bar, 100 µm. (H) Whole-body VO_2_ normalized by body weight of DIO mice treated with saline or rMSA. n = 4 mice per group. (I) Body temperature of DIO mice exposed to cold. n = 6 mice per group. (J) Abundance of thermogenic-related, common adipocyte and inflammation-related mRNAs in BAT from DIO mice treated saline or rMSA. n = 5 mice per group. (K) The protein levels of UCP1 and PGC1α in brown adipose tissues from DIO mice treated with saline or rMSA. n = 3 biological replicates per group. (L) Representative UCP1 immunohistochemical staining in BAT. Scale bar, 100 μm. Data are presented as mean ± s.e.m; \**P* < 0.05, \*\**P* < 0.01, \*\*\**P* < 0.001, \*\*\*\**P* < 0.0001 compared to control group. *ns*, not significant; two-tailed Student’s t-tests (D-F, and J), two-way ANOVA followed by Sidak’s multiple comparisons (B, H, and I).

The reduction in adiposity was accompanied by increased whole-body VO2 (Fig. 4H) and improved maintenance of body temperature during cold exposure (Fig. 4I). BAT from rMSA-treated DIO mice showed increased expression of thermogenic genes and reduced expression of inflammatory markers (Fig. 4J). The protein abundance of PGC1α and UCP1 was also increased, and enhanced UCP1 expression was confirmed by immunohistochemistry (Fig. 4K, L).

The therapeutic effects of rMSA extended beyond the DIO model. In ob/ob mice, rMSA similarly attenuated body-weight gain, reduced fat mass and fat percentage, increased lean mass and lean percentage, lowered liver, and adipose-tissue weights, improved adipose-tissue histology, and reduced hepatic TG content (Fig. S4D–I). In DIO mice, rMSA improved both glucose and insulin tolerance (Fig. S4J, K), and similar improvements were observed in ob/ob mice (Fig. S4L, M).

To determine the contribution of UCP1 to these effects, we treated HFD-fed Ucp1-knockout (UKO) mice with rMSA. Compared with WT mice, the reduction in adiposity induced by rMSA was markedly attenuated in UKO mice, and rMSA failed to increase whole-body VO₂; the histological improvement in BAT was also diminished (Fig. S5). Together, these findings indicate that UCP1 contributes substantially to the anti-adiposity and thermogenic effects of rMSA.

### rMSA promotes thermogenic maturation through Cav1-associated Src– PI3K/AKT–mTORC1 signaling

To define the cellular mechanisms underlying albumin-mediated thermogenesis, we first asked whether rMSA directly regulates thermogenic activity in primary brown adipocytes. Cell viability was largely preserved at rMSA concentrations up to 20 mg/ml, whereas 40 mg/ml significantly reduced viability (Fig. 5A). rMSA increased Ucp1 expression in a concentration-dependent manner, with a maximal response at approximately 10 mg/ml (Fig. 5B), and direct rMSA treatment increased PGC1α and UCP1 protein abundance (Fig. 5D and Fig. S6A). Notably, immobilization of rMSA on agarose beads abolished this effect (Fig. S6B), supporting a direct action of soluble rMSA on brown adipocytes. Because p38/ATF2 signaling is a well-established regulator of thermogenic gene expression in brown adipocytes^34^, we next asked whether this pathway mediated the response to rMSA. rMSA transiently increased p38 and ATF2 phosphorylation (Fig. S6C). However, inhibition of p38 with SB203580 efficiently suppressed ATF2 phosphorylation but had little effect on rMSA-induced PGC1α and UCP1 expression (Fig. S6D). Thus, although rMSA activates p38/ATF2 signaling, this pathway does not appear to be required for the major induction of thermogenic proteins by rMSA.

**Figure 5.**
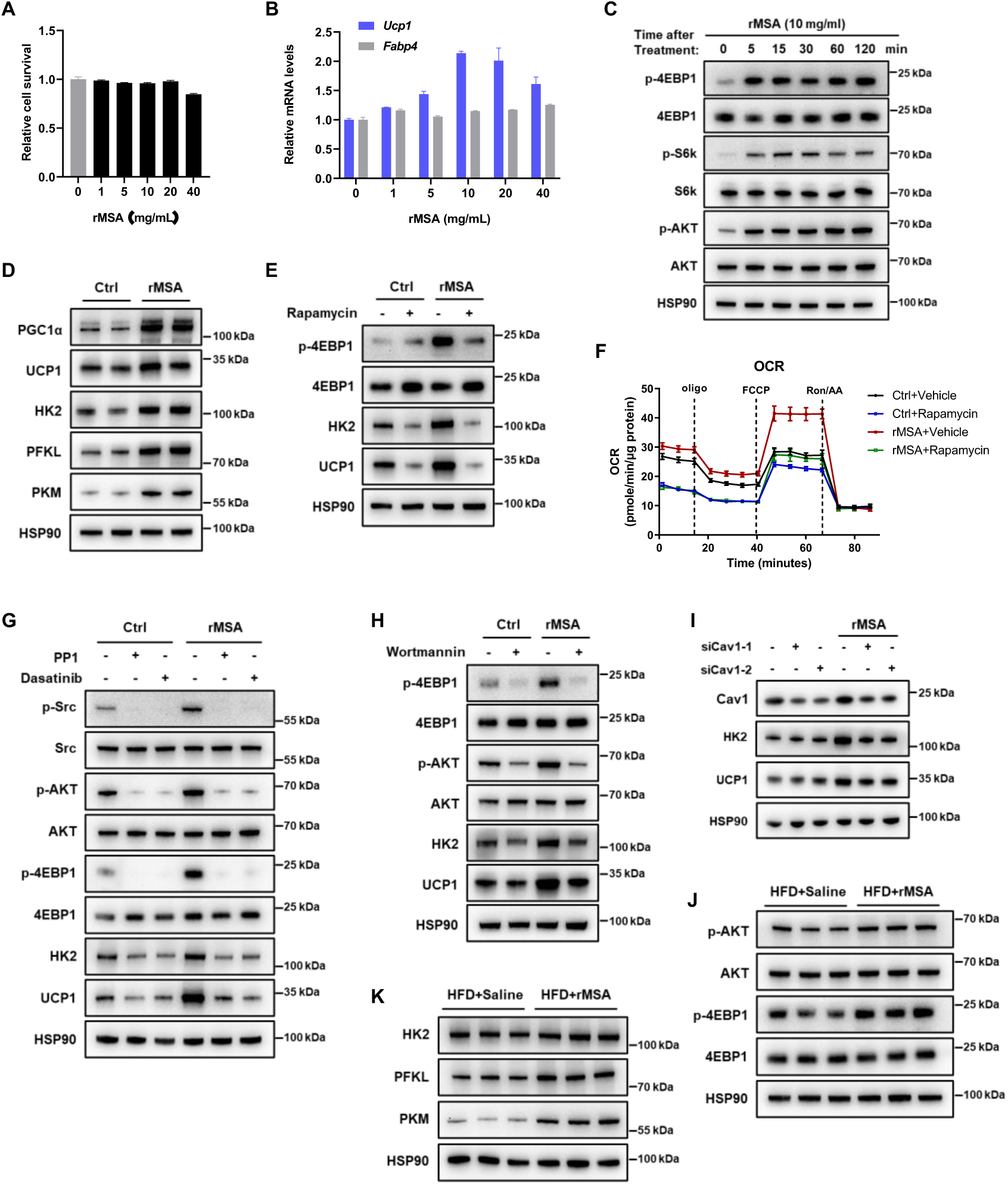
rMSA promotes brown-adipocyte thermogenic maturation through Cav1-associated Src–PI3K/AKT–mTORC1 signaling. (A) CCK-8 viability assay in primary brown adipocytes treated with rMSA. (B) Ucp1 and Fabp4 mRNA abundance after rMSA treatment. (C) Immunoblot analysis of signaling proteins at the indicated times after rMSA treatment. (D) Immunoblots after 48 h rMSA treatment beginning on day 5 of differentiation. (E, F) Effects of rapamycin (100 nM, 30-min pretreatment) on rMSA-induced protein expression (E) and OCR measured by Seahorse mitochondrial stress testing (F) after 48 h treatment. (G, H) Effects of Src inhibition with PP1 (20 μM) or dasatinib (2 μM; G) and PI3K inhibition with wortmannin (1 μM; H), each added 30 min before 48 h rMSA treatment. (I) Immunoblots in primary brown adipocytes transfected with control or Cav1-targeting siRNAs and treated with rMSA for 48 h. (J, K) AKT/4EBP1 phosphorylation (J) and HK2, PFKL and PKM abundance (K) in BAT from DIO mice treated with saline or rMSA; n = 3 biological replicates per group. Data are presented as mean ± s.e.m; \**P* < 0.05, \*\**P* < 0.01, \*\*\**P* < 0.001, \*\*\*\**P* < 0.0001 compared to control group. *ns*, not significant; two-way ANOVA followed by Sidak’s multiple comparisons (F).

Given the established roles of insulin-associated PI3K/AKT signaling and mTORC1 in brown-adipocyte development and thermogenic function^19,35^, we next examined whether this pathway contributes to the effects of rMSA. rMSA rapidly increased AKT phosphorylation and phosphorylation of the mTORC1 downstream targets S6K and 4EBP1 (Fig. 5C), indicating activation of the AKT-mTORC1 signaling pathway. To determine whether this signaling response was associated with thermogenic maturation, we focused on the late stage of brown-adipocyte differentiation. Treatment with rMSA during days 5–6 of differentiation markedly increased PGC1α and UCP1 abundance together with the glycolytic enzymes HK2, PFKL, and PKM (Fig. 5D). Consistent with these protein changes, rMSA also increased glucose consumption and lactate production (Fig. S6F).

We next tested the functional requirement for this signaling pathway. Inhibition of mTORC1 with rapamycin suppressed rMSA-induced 4EBP1 phosphorylation and UCP1 expression (Fig. 5E) and markedly attenuated the increase in cellular oxygen consumption induced by rMSA (Fig. 5F).

Building on prior studies linking albumin–caveolar signaling to Src and PI3K/AKT activation^36–38^, we next asked whether Cav1 and the Src–PI3K/AKT pathway were required for the thermogenic effects of rMSA. Pharmacological inhibition of Src with dasatinib or PP1, or inhibition of PI3K with wortmannin, attenuated rMSA-induced AKT/mTORC1 activation and the associated increases in thermogenic and glycolytic proteins (Fig. 5G, H), supporting a Src– PI3K/AKT–mTORC1 signaling cascade. Importantly, depletion of Cav1 attenuated rMSA-induced HK2 and UCP1 expression (Fig. 5I), indicating that Cav1 is functionally required for the thermogenic and metabolic maturation induced by rMSA. Consistent with UCP1 serving as a major functional effector, rMSA increased respiration in WT brown adipocytes but failed to enhance respiration in UCP1-deficient adipocytes (Fig. S6E).

We next asked whether the signaling changes observed in cultured brown adipocytes were also evident in vivo. BAT from rMSA-treated DIO mice showed increased AKT and 4EBP1 phosphorylation together with increased expression of HK2, PFKL, and PKM (Fig. 5J, K), providing concordant evidence that the pathway identified in vitro is engaged in obese mice. In addition to its effects on mature brown adipocytes, rMSA activated ERK1/2 in brown preadipocytes and increased preadipocyte proliferation, whereas inhibition of either ERK1/2 or mTORC1 attenuated this proliferative response (Fig. S6G–I). Together, these findings identify a Cav1-associated Src–PI3K/AKT–mTORC1 signaling mechanism as a major mediator of rMSA-induced brown-adipocyte thermogenic maturation and further indicate that rMSA promotes brown-preadipocyte proliferation through ERK1/2- and mTORC1-dependent mechanisms.

### Albumin deficiency impairs lipid mobilization during metabolic challenge

Having established that albumin enhances brown-adipocyte thermogenic capacity, we next asked whether it also supports lipid mobilization during increased metabolic demand. Because albumin is the major extracellular carrier of non-esterified fatty acids ^24^, we examined whether albumin deficiency impairs the mobilization of stored lipid under metabolic and adrenergic stimulation.

Under basal conditions, circulating free fatty acid (FFA) and glycerol concentrations were similar between WT and iAKO mice (Fig. 6A, B). In contrast, fasting, cold exposure, and stimulation with the β3-adrenergic agonist CL316243 each elicited markedly smaller increases in circulating FFA and glycerol in iAKO mice (Fig. 6A, B). Consistently, iAKO mice exhibited less hepatic BODIPY-positive lipid accumulation and lower hepatic TG content following these challenges (Fig. 6C–E), supporting impaired mobilization and redistribution of stored lipid.

**Figure 6.**
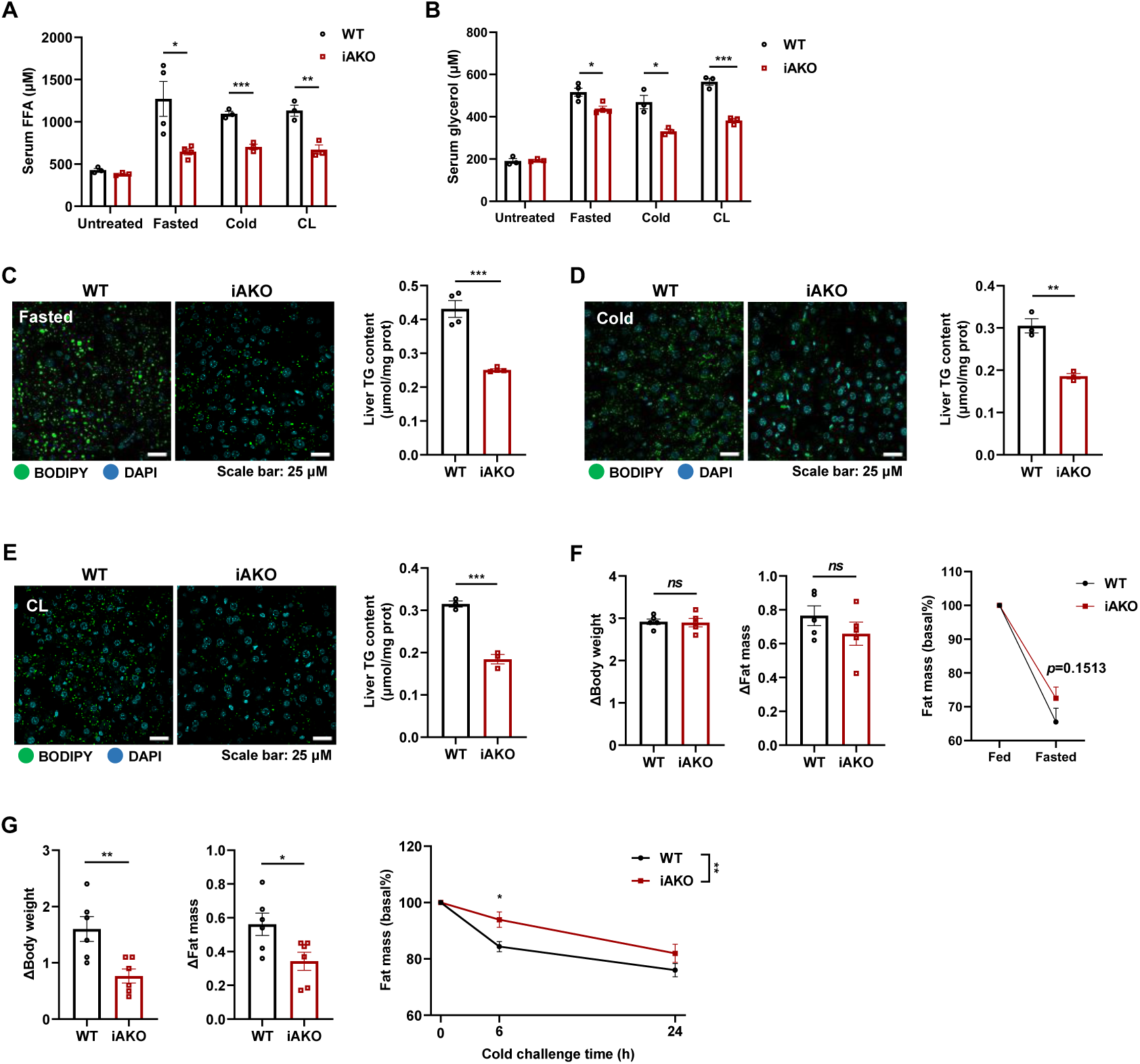
Albumin deficiency impairs lipid mobilization during metabolic challenge. (A) Serum free fatty acid (FFA) concentrations in WT and iAKO mice at baseline and following fasting, cold exposure, or CL316243 (CL) stimulation. n = 3 mice per group; for the fasting condition, n = 4 mice per group. (B) Serum glycerol concentrations in WT and iAKO mice at baseline and following fasting, cold exposure, or CL316243 stimulation. n = 3 mice per group; for the fasting condition, n = 4 mice per group. (C-E) Representative BODIPY staining and quantification of hepatic triglyceride (TG) content in WT and iAKO mice following fasting (C), cold exposure (D), or CL stimulation (E). BODIPY, neutral lipids (green); DAPI, nuclei (blue). Scale bars, 25 μm. Hepatic TG content was normalized to total protein. n = 4 mice per group for fasting and n = 3 mice per group for cold exposure and CL stimulation. (F) Changes in body weight (left) and fat mass (middle), and relative fat mass before and after fasting (right), in WT and iAKO mice. n = 5 mice per group. (G) Changes in body weight (left) and fat mass (middle), and relative fat mass during cold exposure (right), in WT and iAKO mice. Fat mass is expressed relative to the respective baseline value. n = 5 mice per group. Data are presented as mean ± s.e.m; \**P* < 0.05, \*\**P* < 0.01, \*\*\**P* < 0.001, \*\*\*\**P* < 0.0001 compared to control group. *ns*, not significant; two-tailed Student’s t-tests (A-G), two-way ANOVA followed by Sidak’s multiple comparisons (G).

We next asked whether this defect in lipid mobilization translated into altered changes in adiposity during metabolic challenge. Differences in body-weight and fat-mass loss were relatively modest during fasting (Fig. 6F). In contrast, during cold exposure, iAKO mice lost significantly less body weight and fat mass than WT controls (Fig. 6G). Thus, the requirement for albumin became particularly evident under conditions of high thermogenic and adrenergic demand.

To determine whether this defect reflected a broader disturbance in systemic lipid handling, we performed a separate oral lipid challenge. iAKO mice developed prolonged hypertriglyceridemia after olive-oil administration (Fig. S7A), whereas increased TG production did not account for this phenotype (Fig. S7B). Moreover, fatty-acid uptake into iWAT and BAT was comparable between genotypes (Fig. S7C), arguing against enhanced adipose fatty-acid uptake as a major contributor to the excessive fat accumulation observed in albumin-deficient mice.

These findings prompted us to consider whether the fatty-acid burden relative to available albumin, rather than albumin concentration alone, might determine albumin functional capacity. The serum FFA-to-albumin molar ratio was significantly higher in iAKO mice at baseline and remained elevated following fasting, cold exposure, or CL316243 stimulation (Fig. S8A). The ratio was also increased in HFD-fed and ob/ob mice, although the change was less pronounced than that associated with severe albumin deficiency (Fig. S8B). We also considered whether biochemical damage to albumin might further limit its fatty-acid-handling capacity, because obesity is associated with oxidative stress and biochemical modification of circulating albumin^27,39^. Circulating albumin glycation was increased in HFD-fed and ob/ob mice in our study (Fig. S8C). We therefore compared native, glycated, and oxidized rMSA. ANS-based assays, free-thiol measurements, and isothermal titration calorimetry demonstrated marked biochemical changes in GA-rMSA and Ox-rMSA together with impaired fatty-acid binding (Fig. S8D–G). Thus, both the fatty-acid burden relative to available albumin and the biochemical integrity of albumin may determine its effective fatty-acid-handling capacity. Because the defects in albumin-deficient mice were particularly pronounced during cold exposure and β3-adrenergic stimulation, we next tested directly whether the fatty-acid-loading state of albumin regulates β3-adrenergic lipolytic responsiveness.

### Albumin fatty-acid loading regulates β3-adrenergic lipolysis through a Cav1/CD36-associated mechanism

We first asked whether β3-adrenergic lipolytic responsiveness is determined by albumin concentration itself or by the fatty-acid burden relative to available albumin. Across the concentration range tested, changing rMSA concentration alone had little effect on CL316243-stimulated glycerol release from primary brown adipocytes (Fig. 7A). We therefore independently manipulated the two components of the OA:rMSA ratio. When rMSA was held constant at 60 μM, increasing OA from 60 to 180 and 300 μM progressively suppressed CL316243-stimulated glycerol release. Conversely, when OA was held constant at 300 μM, decreasing rMSA from 300 to 100 and 60 μM produced a similar reduction in glycerol release (Fig. 7B). These complementary experiments indicate that β3-adrenergic lipolytic responsiveness is governed by the fatty-acid burden relative to available albumin rather than by either variable alone. Consistent with this relationship, an OA:rMSA molar ratio of 1:1 enhanced CL316243-stimulated glycerol release in both primary white and brown adipocytes, whereas increasing the ratio to 5:1 markedly attenuated this response (Fig. 7C). In contrast, the difference between the 1:1 and 5:1 condition was substantially reduced when lipolysis was stimulated with forskolin (Fig. 7D, E), suggesting that the albumin-sensitive regulatory step lies upstream of direct adenylyl-cyclase activation.

**Figure 7.**
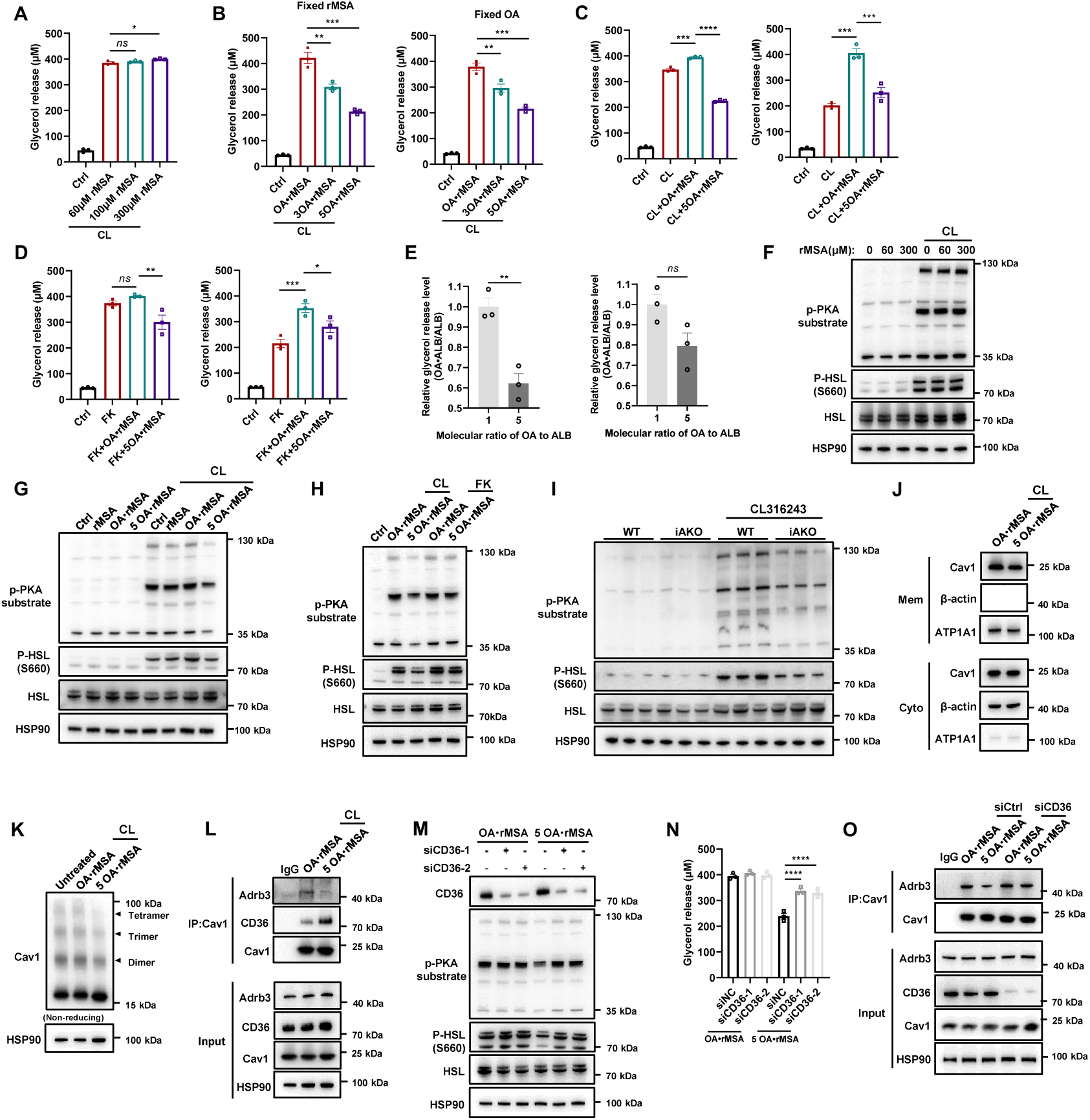
Albumin fatty-acid loading regulates β3-adrenergic lipolysis through a Cav1/CD36-associated mechanism. Unless otherwise indicated, OA:rMSA = 1:1 denotes 300 μM OA plus 300 μM rMSA, and OA:rMSA = 5:1 denotes 300 μM OA plus 60 μM rMSA. (A) Glycerol release from primary brown adipocytes treated for 6 h with the indicated rMSA concentrations ± CL316243 (CL, 10 nM). (B) Glycerol release after 6 h of CL stimulation with fixed rMSA (60 μM) and increasing OA (60, 180 or 300 μM; left), or fixed OA (300 μM) and decreasing rMSA (300, 100 or 60 μM; right). (C, D) Glycerol release from primary white and brown adipocytes treated for 6 h with the indicated OA:rMSA conditions ± CL (C) or forskolin (FK, 10 μM; D). (E) Relative glycerol release calculated from C and D, with the 1:1 condition normalized to 1. (F–H) Immunoblot analysis of lipolytic signaling after 2 h treatment with the indicated rMSA concentrations ± CL (F), OA:rMSA conditions ± CL (G), or OA:rMSA conditions with CL or FK (H). (I) Lipolytic signaling in WT and iAKO BAT under basal or CL-stimulated conditions. n = 3 biological replicates per group. (J) Cav1 in membrane and cytosolic fractions; ATP1A1 and β-actin served as fraction markers. (K) Cav1 oligomerization after DSP crosslinking and non-reducing immunoblotting. (L) Cav1 immunoprecipitation after DSP crosslinking under the indicated OA:rMSA conditions; IgG served as a negative control. (M–O) Effects of CD36 knockdown on lipolytic signaling (M), glycerol release (N), and Cav1-associated Adrb3/CD36 complexes (O). n = 3 biological replicates per group for A–E and N. Data are presented as mean ± s.e.m; \**P* < 0.05, \*\**P* < 0.01, \*\*\**P* < 0.001, \*\*\*\**P* < 0.0001 compared to control group. *ns*, not significant; two-tailed Student’s t-tests (E); one-way ANOVA followed by Dunnett’s multiple-comparisons test (A-D and N).

We next examined whether this loading-dependent effect was reflected in β3-adrenergic signaling. Consistent with the glycerol-release data, varying rMSA concentration alone had little effect on CL316243-induced phosphorylation of PKA substrates and HSL at Ser660 (Fig. 7F). By contrast, the 1:1 OA:rMSA condition potentiated CL316243-induced PKA-substrate and HSL Ser660 phosphorylation, whereas the 5:1 condition markedly attenuated these responses (Fig. 7G). This loading-dependent difference was evident with CL316243 but was largely bypassed when adenylyl cyclase was directly activated with forskolin (Fig. 7H). In vivo, CL316243-induced phosphorylation of PKA substrates and HSL was also reduced in BAT from iAKO mice (Fig. 7I). Together, these findings indicate that the fatty-acid-to-albumin balance regulates β3-adrenergic signaling at a receptor-proximal step upstream of adenylyl cyclase.

Given the reported role of Cav1 in maintaining β3-adrenergic responsiveness, with loss of Cav1 impairing β3-AR signaling and adrenergically stimulated lipolysis^40–42^, we asked whether the fatty-acid-loading state of albumin modulates β3-AR signaling through Cav1-associated membrane organization. Despite the marked difference in β3-adrenergic signaling between the 1:1 and 5:1 OA:rMSA conditions, Cav1 abundance in the membrane fraction was largely unchanged (Fig. 7J), suggesting that altered Cav1 membrane localization alone was unlikely to account for the loading-dependent effect. We therefore examined the higher-order organization of Cav1. Non-reducing immunoblotting revealed loading-dependent differences in Cav1 oligomerization (Fig. 7K). Previous work has shown that fatty acids induce caveolae-dependent CD36 trafficking accompanied by coordinated CAV1 redistribution^43^, prompting us to determine whether albumin fatty-acid loading alters the composition of Cav1-associated membrane complexes. Co-immunoprecipitation revealed stronger Cav1–Adrb3 association under the 1:1 OA:rMSA condition, whereas the higher OA:rMSA ratio (5:1) increased Cav1– CD36 association while concomitantly reducing Cav1–Adrb3 association (Fig. 7L). These findings suggest that high fatty-acid loading shifts the composition of Cav1-associated complexes away from β3-AR and toward CD36. Consistent with this model, CD36 knockdown substantially restored phosphorylation of PKA substrates and HSL at Ser660 under the 5:1 OA:rMSA condition (Fig. 7M). Importantly, CD36 depletion also restored CL316243-stimulated glycerol release under high fatty-acid loading (Fig. 7N), demonstrating functional rescue of lipolysis rather than signaling alone. Moreover, Cav1 immunoprecipitation after CD36 depletion showed restoration of Cav1–Adrb3 association under the high-loading condition (Fig. 7O). Together, these findings indicate that CD36 contributes to high-fatty-acid-loading-induced remodeling of Cav1-associated complexes and is functionally required for the resulting suppression of β3-adrenergic signaling and lipolysis.

These loading experiments therefore indicate that β3-adrenergic responsiveness is determined by the balance between fatty-acid burden and available albumin, with excessive fatty-acid loading suppressing receptor-proximal signaling through a CD36-dependent remodeling of Cav1-associated complexes. We next asked whether biochemical damage that compromises the intrinsic fatty-acid-binding capacity of albumin would produce a comparable functional defect. Consistent with their impaired binding properties (Fig. S8D– G), GA-rMSA and Ox-rMSA were substantially less effective than native rMSA in promoting glycerol release from primary white and brown adipocytes (Fig. S8H). Moreover, native rMSA administration reduced circulating albumin glycation in HFD-fed and ob/ob mice (Fig. S8I), while biochemical comparison of albumin preparations confirmed the relatively preserved integrity of the rMSA used in these studies (Fig. S8J). Together, these findings link preservation of albumin biochemical integrity and fatty-acid-binding capacity to its ability to support adipocyte lipolysis.

Finally, we asked whether preservation of fatty-acid-binding capacity was required for the metabolic effects of albumin in vivo. Whereas native rMSA reduced body weight and adiposity in iAKO, DIO, and ob/ob mice, supplementation with GA-rMSA or Ox-rMSA was substantially less effective across these models (Fig. S9A-I). Biochemical and exposure analyses did not reveal obvious protein aggregation or differences in short-term circulating exposure among the three albumin preparations, although their long-term circulating levels diverged (Fig. S9J, K). Together, these findings suggest that biochemical modification reduces the metabolic efficacy of albumin, although altered long-term disposition may also contribute.

## DISCUSSION

Albumin is classically viewed as an abundant circulating carrier protein, but it also participates in diverse metabolic and signaling processes^32,44^. Despite these established roles, its contribution to the regulation of whole-body energy balance and obesity has remained poorly defined. Here, our findings identify circulating albumin as an active regulator of energy balance and adaptive thermogenesis. Albumin deficiency increased susceptibility to diet-induced adiposity and hepatic lipid accumulation, reduced whole-body energy expenditure and cold tolerance, and impaired stimulus-induced lipid mobilization. Conversely, restoration or administration of rMSA increased thermogenic capacity and alleviated obesity-related phenotypes in multiple mouse models, with UCP1 contributing substantially to its anti-adiposity effects. Mechanistically, our data support two complementary actions of albumin. First, rMSA promotes brown-adipocyte thermogenic maturation through a Cav1-associated Src–PI3K/AKT–mTORC1 mechanism. Second, the balance between fatty-acid burden and available albumin rapidly tunes β3-adrenergic responsiveness through remodeling of Cav1-associated β3-AR/CD36 complexes. Together, these findings extend the conventional view of albumin as a passive fatty-acid carrier and suggest that both its abundance and functional state contribute to metabolic homeostasis.

As albumin is a circulating ligand, identification of the cell-surface receptors and binding proteins that mediate its signaling effects will be important for defining its tissue-specific actions. Previous studies have shown that albumin can engage gp60-dependent caveolar pathways, activate Src kinase, and promote PI3K/AKT signaling^36–38,45–49^. These observations provide a potential framework for our finding that rMSA activates the Src–PI3K/AKT–mTORC1 pathway in brown adipocytes. Rather than acting solely as an acute signaling molecule, our data suggest that albumin can exert a relatively sustained effect on brown-adipocyte function by promoting thermogenic maturation and supporting UCP1-associated thermogenic capacity. In this context, activation of PI3K/AKT/mTORC1 may represent one mechanism through which circulating albumin helps establish or preserve the thermogenic competence of brown adipose tissue.

This longer-term effect is complemented by a more rapid mechanism that becomes apparent during adrenergic stimulation. β-Adrenergic signaling is a major driver of acute thermogenic activation in brown adipocytes^50,51^. Our complementary loading experiments showed that increasing fatty-acid burden at fixed albumin concentration and decreasing albumin availability at fixed fatty-acid concentration both attenuated β3-adrenergically stimulated lipolysis. These findings indicate that adrenergic responsiveness is governed by the balance between fatty-acid burden and available albumin rather than by either variable alone. Mechanistically, relatively low fatty-acid loading favored Cav1 association with β3-AR and robust downstream PKA/HSL signaling, whereas high fatty-acid loading shifted Cav1-associated complexes toward CD36 and away from β3-AR. Importantly, CD36 depletion not only restored PKA/HSL signaling but also rescued glycerol release and Cav1–β3-AR association under high-loading conditions, supporting a functional role for CD36 in this receptor-proximal remodeling process. Thus, albumin may coordinate thermogenesis across distinct time scales: its signaling actions contribute to the development and maintenance of thermogenic capacity, whereas the relationship between fatty-acid burden and albumin availability rapidly tunes adrenergic responsiveness according to the prevailing lipid environment. This model also provides a physiological link between albumin’s classical function as a fatty-acid carrier and its newly identified role in regulating energy expenditure. Given that albumin is predominantly produced by the liver and acts directly on brown adipocytes in vitro, our findings further raise the possibility that circulating albumin functions as a liver-derived metabolic signal linking hepatic secretory function to brown-adipose thermogenesis. In this context, albumin may exhibit hepatokine-like properties and contribute to interorgan coordination of systemic energy homeostasis. Nevertheless, the receptor responsible for albumin-induced Src–PI3K/AKT–mTORC1 activation remains unresolved. Although CD36 depletion supports a causal role for CD36 in the high-fatty-acid-loading-induced suppression of β3-adrenergic responsiveness, whether the redistribution of Cav1 between β3-AR- and CD36-associated complexes reflects direct competition, altered trafficking, or broader remodeling of caveolar organization remains to be determined.

The health benefits of albumin are associated with its quality^31^. At present, the clinical utilization of HSA is mainly derived from donor blood^52^, and its biochemical status may vary depending on donor background, storage, and accumulated molecular modifications. Notably, a recent randomized, double-blind phase 3 trial in patients with cirrhotic ascites demonstrated clinical equivalence of recombinant HSA to plasma-derived HSA in restoring serum albumin and non-inferiority in improving ascites, supporting the clinical feasibility of well-defined recombinant albumin products^53^. Previous studies have linked damaged or modified albumin to metabolic diseases, including T2DM and obesity^25,39,54^. Our findings further suggest that such biochemical integrity is functionally important: glycation and oxidation markedly impaired the fatty-acid-binding capacity of albumin and substantially reduced its metabolic efficacy in vivo. This concept is supported by recent findings in an independent biological context. A recent *Nature* study showed that albumin protects its bound fatty acids from oxidation and thereby preserves their antifungal activity against Mucorales; importantly, albumin glycation caused dissociation and oxidation of bound fatty acids and markedly impaired this protective function^55^. Thus, the biochemical quality of albumin can directly determine the stability and biological activity of its associated lipid cargo. A similar principle has been observed for other metabolically important circulating proteins. For example, glycation of apolipoprotein A-I alters its structure, reduces lipid-binding and cholesterol-efflux capacity, and impairs its metabolic functions^56^. Together, these observations emphasize that the biological activity of abundant circulating carrier proteins depends not only on their concentration, but also on preservation of their structural and ligand-binding integrity. The rMSA used in our studies has previously been characterized as ultrapure, young, and minimally damaged^31^, which may contribute to its preserved metabolic activity and supports the potential value of well-defined recombinant albumin products. However, the glycated and oxidized rMSA preparations used here represent experimentally defined forms of damage and may not fully recapitulate the heterogeneous modifications present in circulating albumin in metabolic disease. The reduced metabolic efficacy of modified rMSA was not accompanied by obvious high-molecular-weight aggregation or degradation, and the three preparations showed comparable short-term circulating exposure after intravenous administration. Their circulating levels nevertheless diverged during the later phase, indicating that altered long-term disposition may also contribute to differences in in vivo efficacy. Accordingly, impaired fatty-acid binding is likely to represent one mechanistic consequence of albumin damage, but cannot be considered the sole explanation for the reduced metabolic activity of glycated or oxidized albumin. Potential effects of biochemical modification on cellular uptake, receptor engagement, tissue distribution, or downstream signaling also remain to be defined.

Collectively, our findings provide a framework for understanding albumin as a dynamic regulator of metabolic homeostasis, in which albumin abundance, fatty-acid loading state, and biochemical integrity jointly influence brown-adipose thermogenic capacity and adrenergic responsiveness. These findings further highlight preservation of albumin quality and ligand-handling function as important considerations for future therapeutic development.

## METHODS

### Mice models

C57BL/6J (WT) mice were purchased from Charles River. Constitutive albumin-knockout mice (C57BL/6J-Albem8Mvw/MvwJ; AKO) were purchased from The Jackson Laboratory. AKO litters were bred as homozygous males (*Alb*^−/−^) crossed with heterozygous females (*Alb*^+/−^), as recommended by The Jackson Laboratory. ob/ob (B6.V-Lepob) male mice (SPF grade) were purchased from HFK Bioscience (Beijing, China). *Alb^fl/fl^* mice were purchased from GemPharmatech and used to generate inducible albumin-knockout (iAKO) mice as described below. Ucp1-knockout (*Ucp1*^−/−^; UKO) male mice were purchased from Shanghai model organisms Center and used to assess the contribution of UCP1 in the metabolic effects of rMSA.

All mice were weighed and allocated to experimental groups according to body weight using a random-number table. Mice were housed in pathogen-free barrier facilities under a 12-h light/dark cycle in individually ventilated cages (maximum six mice per cage) at 22– 26 °C, with sterile pellet food and water available ad libitum unless otherwise indicated. Animals were monitored daily. The animal facility is accredited by AAALAC International, and all procedures were approved by the Institutional Animal Care and Use Committee of Tsinghua University (approval numbers 18-LYZ2 and 21-LYZ1) in accordance with the Guide for the Care and Use of Laboratory Animals.

Mice were acclimatized for 1 week after arrival. Unless otherwise indicated, male mice 6– 8 weeks of age were used. For diet-induced obesity (DIO), 8-week-old male mice were fed a high-fat diet (HFD; Research Diets, D12492) for 8 weeks before the indicated interventions. HFD feeding was continued during subsequent treatment experiments unless otherwise specified.

### Generation of inducible albumin-knockout mice

To generate inducible albumin-knockout (iAKO) mice, *Alb^fl/fl^*mice were administered AAV2/8-hTBG-iCre-eGFP-WPRE-pA (Taitool Bioscience) via tail-vein injection. The viral stock had a titer of 1.2 × 10^13^ viral genomes (vg)/mL and was diluted to 1 × 10^12^ vg/mL for administration. Each mouse received 1 × 10^11^ vg of AAV. Wild-type C57BL/6J mice on the same genetic background as the *Alb^fl/fl^*mice received the same AAV preparation at the same dose and by the same route and served as WT controls. Following AAV administration, circulating albumin levels were measured to confirm albumin depletion before HFD challenge.

### Immobilized rMSA treatment

For immobilized rMSA treatment, NHS-activated agarose (Sangon Biotech, C600024) was used to bind rMSA following the manufacturer’s instruction. Briefly, the agarose was washed and equilibrated with 1 mM HCl and sodium bicarbonate buffer (pH = 8.0), respectively. Next, rMSA was co-incubated with the agarose or the agarose alone overnight at 4 °C in sodium bicarbonate buffer (pH = 8.0). The samples were then blocked with 0.1 M Tris-HCl buffer (pH = 8.5). Free rMSA, immobilized rMSA, or empty agarose were added to Transwell chambers with 4 μm pore membranes positioned above the cells. After 24 hours of treatment, cells were subjected to western blot analysis to assess thermogenesis.

### rMSA and modified-albumin administration

The young, undamaged, and ultrapure rMSA and treatments used in this study were the same as previously described ^31,32^. Unless otherwise indicated, mice in the rMSA-treated groups received rMSA intravenously at 1.5 mg/g body weight once every 3 weeks for a total of three administrations, whereas control mice received an equal volume of saline according to the same schedule. For experiments comparing native and chemically modified albumin, mice received native rMSA, glycated rMSA (GA-rMSA), or oxidized rMSA (Ox-rMSA) at the same dose and administration schedule.

Circulating albumin was assessed using different methods according to the experimental model. In iAKO mice and rMSA-treated iAKO mice, serum was collected and albumin abundance was assessed by immunoblotting. In DIO, ob/ob, and other animal experiments, serum albumin concentrations were measured using the bromocresol green (BCG) method.

### Metabolic cages

The energy budget of mice was measured using the TSE PhenoMaster metabolic cages housed within environment-controlled rodent incubators at Tsinghua University. Mice were singly housed. Before data collection, mice were acclimatized in metabolic chambers for 24 h. Mice had free access to food and water. Physical activity, food intake, CO_2_, and O_2_ levels were collected about every 27 minutes over the duration of the experiment. Whole-body VO₂ and energy expenditure were calculated from the PhenoMaster output and expressed relative to body weight as indicated in the corresponding figures.

### Mouse body composition analysis

For assessing metabolic parameters, body weight and body compositions (lean and fat masses) were measured by quantitative magnetic resonance (MR) using Echo MRI-100 Body Composition Analyzer (EchoMRI).

### Histological, immunohistochemistry, and lipid staining

Adipose and liver were fixed in 4% paraformaldehyde solution overnight at room temperature and embedded in paraffin before sectioning. Sections were successively stained with hematoxylin and eosin and captured by a Digital Pathology Section Scanner (KFBIO, KF-PRO-120). Adipocyte areas were quantified by Image-Pro Plus 6.0 (Media Cybernetics).

For immunohistochemical (IHC) staining of UCP1, sections were incubated with anti-Rabbit UCP1 (CST) at a 1:200 dilution overnight and detected according to the general manufacturer’s instructions.

For Lipid staining, liver tissues were embedded in optimal cutting temperature (OCT) compound and prepared as frozen sections. Sections were stained with BODIPY 493/503 to visualize neutral lipid accumulation, washed with PBS, and mounted for fluorescence imaging. BODIPY-positive lipid accumulation was captured using a fluorescence microscope (FV3000, Olympus).

### Hepatic triglyceride quantification

For the detection of triglycerides in the liver, 50 mg of tissue was washed with PBS, minced, and homogenized in 500 µl lysis buffer. After incubation at room temperature for 10 min and centrifugation (2000 rpm, 5 min), the concentration of triglycerides in the supernatant was quantified using a tissue triglyceride content assay kit (E1013, Applygen).

### Glucose, insulin, and pyruvate tolerance tests

For glucose tolerance tests (GTTs), mice were fasted for 14-16 h and injected intraperitoneally with glucose (2 mg/g body weight). Blood glucose was measured at 0, 15, 30, 60, and 120 min using an ACCU-CHEK glucometer (Roche).

For insulin tolerance tests (ITTs), mice were fasted for 4 h and injected intraperitoneally with insulin (0.5 U/kg body weight). Blood glucose was measured at 0, 15, 30, and 60 min using the same glucometer.

For pyruvate tolerance tests, mice maintained on normal diet were fasted 14-16 h before intraperitoneal pyruvate (2 mg/g body weight) administration, and blood glucose was measured serially to assess the glycemic response.

### Oral lipid tolerance assay

Mice were subjected to a 6-hour fasting period before undergoing the oral lipid tolerance assay. A volume of 200 μl olive oil was administered to the mice via oral gavage. Blood samples were collected from the tail vein immediately before the gavage and at various time points thereafter. Serum was then isolated to measure TG levels using a liquid sample triglyceride content assay kit (E1003, Applygen).

### Cold challenge and body temperature measurement

Experiments with cold challenges were carried out in frigid settings. Mice were housed singly in pre-chilled cages at a temperature of 4 °C. Mice had free access to water and pre-chilled food. A digital thermometer and a rectal thermocouple probe (TH-212, HICHANCE) were used to take rectal core body temperatures every two hours. If a mouse’s core body temperature dropped below 20 °C, it was sacrificed, and the death was recorded as an event for the survival analysis.

### Western blotting

Western blotting was performed as we described previously. Briefly, tissues or cells were lysed with Cell Lysis Buffer for Western or IP (Beyotime, Cat# P0013) on ice. Equal amounts of proteins were separated by SDS-PAGE using 4–12% precast gels (LABLEAD) and transferred to polyvinylidene difluoride membranes (Millipore). The membranes were blocked with 5% skimmed milk and incubated with the indicated primary antibodies overnight at 4 °C. Details of antibodies used in the study are provided in Table S1. Proteins were detected using horseradish peroxidase (HRP)-conjugated secondary antibodies (CST) and enhanced chemiluminescence detection reagents (sc-2048, Santa Cruz), and signals were captured with a CCD scanner (4600SF, Tanon).

### Quantitative real-time reverse transcription PCR

Total RNA of tissues was extracted using the TRIzol reagent (Invitrogen), and a First Strand cDNA Synthesis Kit (Fermentas, Thermo Fisher Scientific) was used for reverse transcription according to the manufacturer’s protocol. Quantitative real-time PCR (qRT-PCR) was performed using gene-specific primers on an Mx3000P system (Stratagene). Relative quantification of the gene expression was calculated using the 2^−ΔΔCT^ method and was presented as fold change normalized to that of β-actin. The sequences of primers used in this study were listed in Table S2.

### Cell culture

Primary brown preadipocytes were obtained from SVF of BAT that isolated from 7-day old pups. Post-confluent cells were differentiated with adipogenic cocktail (1 μM rosiglitazone, 0.5 mM IBMX, 5 μM dexamethasone, 0.114 μg/ml insulin, 50 nM T_3_ and 125 μM indomethacin). Cells were re-fed every 48 hours with maintenance cocktail (1 μM rosiglitazone, 50 nM T_3_ and 0.5 μg/ml insulin). Adipocytes were fully differentiated 6 days after initial exposure to the adipogenic cocktail. For primary white adipocyte cultures, SVF cells were isolated from the iWAT of 6-week-old male C57BL/6J mice. Adipocyte differentiation was induced using the same protocol as for primary brown adipocytes, except that 10 μg/mL insulin was maintained throughout differentiation and T3 was omitted. For experiments examining thermogenic maturation, primary brown adipocytes were treated beginning on day 5 of differentiation and maintained under the indicated conditions for 48 h. Where indicated, cells were pretreated for 30 min with rapamycin (100 nM), the Src inhibitors PP1 (20 μM) or dasatinib (2 μM), or the PI3K inhibitor wortmannin (1 μM), followed by rMSA treatment for 48 h.

### In vivo lipid mobilization challenges

To assess lipid mobilization under different metabolic challenges, circulating free fatty acids (FFAs) and glycerol were measured under basal conditions and following fasting, acute cold exposure, or β3-adrenergic stimulation.

For the fasting challenge, mice were fasted overnight for 16 h with free access to water. Blood samples were collected before and after the fasting period. For the cold challenge, mice were individually housed at 4 °C for 6 h with free access to food and water, and blood samples were collected at baseline and after 6 h of cold exposure. For body weight and body composition measurements, mice were subjected to either fasting or cold exposure for 24 h beginning at 12:00, with measurements performed immediately before and after the challenge. Water was provided ad libitum during fasting, whereas food and water were provided ad libitum during cold exposure.

For β3-adrenergic stimulation, mice received CL316243 (Selleck, E1049) at 1 mg/kg by i.p. injection. Blood samples were collected immediately before and 3 h after CL316243 administration. For analysis of acute β3-adrenergic signaling in BAT, mice were euthanized 120 min after CL316243 administration, and interscapular BAT was rapidly dissected, snap-frozen, and subsequently analyzed by immunoblotting for PKA-substrate and HSL phosphorylation.

### Serum FFA and glycerol measurements

Blood samples were collected and centrifuged to obtain serum. Circulating FFA concentrations were measured using the LabAssay™ NEFA (FFA) kit (FUJIFILM Wako Pure Chemical Corporation, Cat# 299-94301), and glycerol concentrations were determined using the Liquid Sample Glycerol Content Assay Kit (Applygen Technologies Inc., Cat# E1002), according to the manufacturers’ instructions. Absorbance was measured at 550 nm using a microplate reader, and concentrations were calculated from the corresponding standard curves.

### Triglyceride production assay

To assess endogenous triglyceride production, mice were fasted 8 h and then given an intraperitoneal injection of poloxamer 407 (HY-D1005, MCE). One hour later, triglyceride levels were measured from the serum and the mice were given a 200-μl oral olive oil bolus. Two hours later, serum triglyceride levels were measured again. Serum was then isolated to measure TG levels using a liquid sample triglyceride content assay kit (E1003, Applygen).

### In vivo fatty acid uptake assay

Tissue fatty-acid uptake was assessed using BODIPY FL C16 (Invitrogen, D3821). Mice were fasted overnight before the experiment and received BODIPY FL C16 (10 mg/kg) by oral gavage. At 30 min after tracer administration, mice were euthanized, and iWAT and BAT were rapidly collected and weighed. Tissues were homogenized in methyl tert-butyl ether (MTBE) to extract lipids. Homogenates were centrifuged at 12,000 × g for 10 min, and fluorescence in the resulting lipid extracts was measured using a microplate reader at excitation/emission wavelengths of 505/512 nm. Fatty-acid uptake was normalized to tissue weight and expressed relative to the corresponding control group.

### ANS fluorescence assay

The fatty-acid-binding properties of native recombinant mouse serum albumin (rMSA), glycated rMSA (GA-rMSA), and oxidized rMSA (Ox-rMSA) were assessed using a 1-anilino-8-naphthalene sulfonic acid (ANS) fluorescence displacement assay according to a previously established protocol with modifications. Briefly, native rMSA, GA-rMSA, or Ox-rMSA (2.5 μg in 100 μL PBS) was added to black 96-well flat-bottom microplates, followed by the addition of 50 μL ANS solution in PBS to achieve final ANS concentrations of 0, 1.25, 2.5, 5, and 10 μM. Samples were incubated for 15 min at room temperature in the dark. For the displacement assay, oleic acid (OA) was added at increasing concentrations (0– 10 μM) to the preformed ANS–albumin complexes. Samples without OA were included to determine the initial fluorescence intensity. After an additional 15-min incubation at room temperature in the dark, fluorescence was measured using a microplate reader (Enspire, PerkinElmer) with excitation and emission wavelengths of 360 and 460 nm, respectively. Fluorescence quenching, indicative of ANS displacement by OA, was recorded as the percentage change in fluorescence intensity (Δ*F*%) using the formula: %Δ*F* quenching = [(*F*_0_ − *F*_1_)/*F*_0_] × 100, where *F*_0_ is the fluorescence in the absence of OA and *F*_1_ is the fluorescence at each tested concentration of OA.

### Measurement of free thiol content

The free-thiol content of native rMSA, GA-rMSA, and Ox-rMSA was determined using 5,5′-dithiobis-(2-nitrobenzoic acid) (DTNB; Ellman’s reagent). Albumin samples were diluted tenfold in 20 mM phosphate buffer containing 1 mM EDTA (pH 7.4). Diluted samples (50 μL) were mixed with 150 μL of 2 mM DTNB working solution and incubated for 5 min at room temperature in the dark. Absorbance was measured at 412 nm using a microplate reader. The free-thiol concentration was calculated from an L-cysteine standard curve (0– 0.08 mM), corrected for sample and reaction-volume dilution, and normalized to the albumin mass.

### In vitro lipolysis assay

Fully differentiated primary adipocytes were incubated in serum-free medium and treated with PBS (vehicle control), rMSA with different oleic acid (OA) loading ratios, glycated rMSA (GA-rMSA), or oxidized rMSA (Ox-rMSA), in the presence or absence of CL316243. After 6 h of treatment, the culture medium was collected, and glycerol released into the medium was quantified using a Glycerol Assay Kit (E1002, Applygen) according to the manufacturer’s instructions.

### Preparation and biochemical characterization of glycated and oxidized rMSA

Briefly, glycated rMSA (GA-rMSA) was prepared by incubating native rMSA in PBS containing 5 mM glucose at 37 °C for 72 h under 95% O₂/5% CO₂ (pH 7.3–7.4)^57^. Oxidized rMSA (Ox-rMSA) was prepared by treating rMSA with a 50 - fold molar excess of L-cysteine/cystine (80 ml of 0.06 mM rMSA mixed with 72 ml of 3 mM cysteine and 8 ml of 3 mM cystine) in 0.1 M sodium carbonate/bicarbonate buffer (pH 10) at 37 °C for 48 h^58^. Both reaction mixtures were then purified from low-molecular-mass components (excess glucose or cysteine/cystine) by ultrafiltration through a 3000-Da membrane (Millipore) at 4 °C. The purified products were stored at 4 °C for subsequent cellular and animal experiments.

Glycated albumin levels were measured using a glycated albumin assay kit (BDH) based on the enzymatic peroxidase method according to the manufacturer’s instructions. Briefly, serum albumin was selectively hydrolyzed by an albumin-specific protease to release glycated amino acids. The resulting glycated amino acids were subsequently oxidized by a specific glycated amino acid oxidase, generating glucosone, amino acids, and hydrogen peroxide. In the presence of peroxidase, hydrogen peroxide reacted with 4-aminoantipyrine (4-AAP) and HTIB through a Trinder reaction to generate a red quinoneimine chromogen. Glycated albumin concentrations were determined from the change in absorbance according to the manufacturer’s protocol.

### Isothermal titration calorimetry

The binding of oleic acid (OA) to native, glycated, and oxidized recombinant mouse serum albumin (rMSA, GA-rMSA, and Ox-rMSA, respectively) was analyzed by isothermal titration calorimetry (ITC). OA was dissolved in ethanol to prepare a 10 mM stock solution and subsequently diluted to the working concentration in assay buffer. Albumin samples were prepared at approximately 20 μM, and the solvent composition was matched between the protein and OA solutions to minimize heat arising from solvent dilution. After thermal equilibration, OA was sequentially titrated into the albumin solution. Control titrations of OA into assay buffer were performed under the same conditions to correct for the heat of dilution. The heat associated with each injection was integrated, and the resulting binding isotherms were fitted using a one-set-of-sites model in the MicroCal PEAQ-ITC Analysis Software. The apparent dissociation constant (Kd,app) and apparent binding stoichiometry (N) obtained from the fitted isotherms were used to compare the OA-binding properties of native, glycated, and oxidized rMSA.

### Loading of albumin with OA

Oleic acid (Maclin, 112-80-1) was saponified with an equimolar amount of NaOH to prepare a 10 mM sodium oleate solution. Briefly, 28.25 mg OA was dissolved in 1 mL ethanol, mixed with 100 μL of 1 M NaOH, and the solution was brought to a final volume of 10 mL. The rMSA solution was pre-equilibrated at 37°C, and the saponified OA solution was subsequently mixed with rMSA at the indicated OA:rMSA molar ratios. For the standard 1:1 and 5:1 condition used in subsequent signaling experiments, OA was maintained at 300 μM and rMSA was used at 300 or 60 μM, respectively.

### Glucose utilization and lactate production in adipocytes

Glucose and lactate concentrations in culture supernatants were measured using commercially available glucose and lactate assay kits according to the manufacturers’ instructions (Beyotime, S0202 and S0208).

For glucose measurement, 5 μl of sample was mixed with 185 μl of Glucose Assay Reagent, heated at 95 °C for 8 min, and subsequently cooled to 4 °C. Aliquots of the reaction mixture were transferred to 96-well plates, and absorbance was measured at 630 nm using a microplate reader. Glucose concentrations were calculated from a glucose standard curve.

For lactate measurement, a WST-8-based working solution containing Lactate Assay Buffer, Enzyme Solution, Substrate, and WST-8 was prepared according to the manufacturer’s protocol. Samples (50 μl) were mixed with an equal volume of the WST-8 working solution and incubated at 37 °C for 30 min in the dark. Absorbance was measured at 450 nm using a microplate reader, and lactate concentrations were determined from an L-lactate standard curve.

### EdU cell proliferation assay

Cell proliferation was assessed using a commercial EdU incorporation assay kit according to the manufacturer’s instructions (Beyotime, C0075). Cells were incubated with 5 μM EdU for 24 h and subsequently collected by trypsinization. Cells were fixed with 4% paraformaldehyde for 30 min at room temperature and permeabilized with 0.2% Triton X-100 in PBS for 10 min. EdU incorporation was detected using the Click reaction mixture for 30 min at room temperature in the dark, followed by nuclear staining with Hoechst 33342 for 5 min. After washing and filtration, cells were analyzed by flow cytometry, and the percentage of EdU-positive cells was used as an index of cell proliferation.

### Cell viability assay

Cell viability was assessed using a commercial Cell Counting Kit-8 (CCK-8) assay according to the manufacturer’s instructions (Beyotime, C0037). Briefly, CCK-8 reagent was diluted 1:9 with serum-free culture medium, and 100 μl of the resulting working solution was added to each well. Cells were incubated at 37 °C for 1–4 h, after which absorbance was measured at 450 nm using a microplate reader. Cell viability was calculated based on the absorbance values relative to the corresponding control group.

### Membrane protein extraction

Membrane and cytosolic proteins were separated from adipocytes using a membrane protein extraction kit (Beyotime, P0033) according to the manufacturer’s instructions. Equal amounts of protein from the membrane and cytosolic fractions were subjected to western blot analysis. ATP1A1 and β-actin were used as markers for the membrane and cytosolic fractions, respectively. The resulting fractions were used to assess Cav1 membrane partitioning.

### DSP crosslinking and Cav1 oligomerization

For stabilization of protein complexes, adipocytes were treated with 1 mM DSP (Selleck, S0626) in PBS for 30 min at room temperature before cell lysis. The crosslinking reaction was quenched with 20 mM glycine in PBS for 15 min at room temperature, after which the cells were washed with PBS and subjected to cell lysis. DSP-crosslinked samples used for Cav1 oligomerization analysis were prepared with non-reducing protein-loading buffer (Beyotime, P0016P) and subjected to SDS-PAGE followed by western blotting for Cav1. DSP-crosslinked cell lysates were also used for co-immunoprecipitation experiments as described below.

### Co-immunoprecipitation

DSP-crosslinked cells were lysed, and equal amounts of protein were incubated with the indicated immunoprecipitating antibodies overnight at 4 °C. Immune complexes were captured using protein A/G beads, washed, and eluted for western blot analysis. An equivalent amount of IgG was used as a negative control. Input lysates and immunoprecipitated samples were analyzed by western blotting with the indicated antibodies. Details of the antibodies used for immunoprecipitation and western blotting are provided in Table S1.

### siRNA-mediated Cav1 or CD36 knockdown

For Cav1 or CD36 knockdown, primary brown adipocytes were transfected with siRNAs targeting Cav1 or Cd36, or with the corresponding negative-control siRNA, using a transfection reagent (GenePharma) according to the manufacturer’s instructions. At 48 h after transfection, cells were subjected to the indicated treatments. For Cav1 knockdown experiments, cells were treated with rMSA for 48 h, and knockdown efficiency together with the expression of thermogenic and metabolic proteins was assessed by western blotting. For CD36 knockdown experiments, cells were treated with the indicated OA:rMSA conditions in the presence of CL316243, and knockdown efficiency and downstream signaling responses were assessed by western blotting. Where indicated, glycerol release was quantified as described above, and DSP crosslinking followed by Cav1 immunoprecipitation was performed as described below. The siRNA sequences used in this study are provided in Table S3.

### Seahorse

Cellular OCR was measured using a XFe96 Analyzer (Agilent Technologies). Primary adipocytes were plated at 5000 cells/well in XF 96-well microplates (Agilent Technologies) and differentiated as described above. On day 5 of differentiation, cells were pretreated with vehicle or rapamycin (100 nM) for 30 min, followed by treatment with vehicle or rMSA (10 mg/ml) for 48 h. Cells were then incubated in pre-warmed assay medium before the assay according to the manufacturer’s instructions (Agilent Seahorse XF Cell Mito Stress Test Kit). For mitochondrial stress test, cells were subjected to oligomycin (2 μM), FCCP (2 μM), and rotenone/antimycin A (1 μM) injection sequentially. At the beginning of the assay, the medium was changed to unbuffered, glucose-free DMEM (Sigma-Aldrich, D5030, pH 7.35 at 37°C) supplemented with 2 mM glutamine. OCR values were normalized to protein content.

### Non-reducing Coomassie staining

Native rMSA, GA-rMSA, and Ox-rMSA were mixed with non-reducing loading buffer without heating and separated on a 4-12% SDS-PAGE gel. Gels were stained with Coomassie Brilliant Blue (Beyotime, P0017) overnight. The assay was used to assess gross high-molecular-weight aggregation and degradation and was not intended to exclude subtle conformational differences.

### Circulating exposure of modified albumin

To compare the circulating exposure of native and modified albumin preparations, AKO mice received a single intravenous injection of rMSA, GA-rMSA, or Ox-rMSA at 1.5 mg/g body weight. Serum samples were collected during the short-term period (1,2,6,12,24, and 48 h) and long-term period (7,14, and 21 days) after injection, and serum albumin concentrations were quantified by Mouse albumin ELISA kit (Elabscience, E-EL-M3032). Before the circulating-exposure study, assay recovery was validated for native rMSA, GA-rMSA and Ox-rMSA at the working dilution. All three preparations showed comparable recovery exceeding 90%.

### RNA sequencing and data analysis

Six mouse BAT samples were included in the transcriptomic analysis, comprising three biological replicates each from the HFD_AKO and HFD_WT groups. Transcriptome sequencing was performed by BGI Wuhan Co., Ltd. RNA integrity and concentration were assessed using an Agilent 2200 TapeStation, and RNA samples with an RNA integrity number (RIN) >8.5 were used for cDNA library preparation. Ribosomal RNA was depleted by hybridization capture, followed by single-end 50-nt sequencing on the BGISEQ-500 platform, generating more than 20 million reads per sample. Sequencing reads were quality-filtered using Trimmomatic v0.33. To generate the PCA, KEGG, and GSEA figures presented in this study, clean reads were quantified against the Ensembl release 102 mouse GRCm38 reference transcriptome using kallisto v0.46.2, and transcript-level counts were aggregated to the gene level. Differential expression analysis was performed using PyDESeq2 v0.5.4, an implementation of the DESeq2 statistical framework, with the contrast AKO versus WT. Genes with a Benjamini–Hochberg-adjusted P value (padj) <0.05 and an absolute log2 fold change (log2FC) ≥0.585 were considered differentially expressed. KEGG over-representation analysis was performed separately for the two gene sets using a hypergeometric test followed by Benjamini–Hochberg correction. For GSEA, all genes were preranked according to the DESeq2 Wald statistic, and normalized enrichment scores (NES), nominal P values, and false discovery rate (FDR) q values were calculated using 2,000 gene-set permutations. All PCA, KEGG, and GSEA figures shown in this study were generated using the reanalysis workflow described above.

### Quantification and statistical analysis

Age- and weight-matched mice were randomly assigned to the experiments. The number of animals used in each experiment is provided in the respective figure legends. No animals were excluded from the statistical analyses, and the investigators were not blinded during the studies. Except for the animal studies and RNAseq, which were conducted once, each experiment was repeated at least three times, yielding consistent results. EThe numbers of biological replicates and animals used in each experiment are specified in the corresponding figure legends. Results are presented as means ± s.e.m. Comparisons of only two groups were performed by Student’s t-test (two-tailed). Statistical analysis for multiple group comparisons was performed by one-way analysis of variance (ANOVA) with Dunnett’s test or two-way ANOVA with Sidak’s test. All statistical tests were two-sided, and all *P* values less than 0.05 were considered statistically significant, with \**P* < 0.05; \*\**P* < 0.01; \*\*\**P* < 0.001; \*\*\*\**P* < 0.0001. Statistical analyses and graphical presentations were performed using GraphPad Prism (GraphPad Software, San Diego, CA, USA). Statistical assays performed are specified in figure legends.

## Data availability

All data supporting the findings of this study are available within the article and its Supplementary Information or from the corresponding author upon reasonable request. The RNA-sequencing data generated in this study have been deposited in the NCBI Gene Expression Omnibus (GEO) under accession code GSE348868.

## Acknowledgements

We thank all the staff members of Luo laboratory for their technical support and critical suggestions.

## Author contributions

YL conceived and coordinated the project. YL and ZL designed the experiments, interpreted the data, and wrote the manuscript. ZL performed most of the experiments. YJ performed a part of the experiments. HL, XD, EL, JZ, and YF assisted with the experiments and helped to analyze the data and revise the manuscript. All authors read and approved the final manuscript.

## LLM disclosure

OpenAI ChatGPT was used to assist with manuscript language refinement, structural organization, and discussion of statistical presentation. All scientific interpretations and final decisions were independently reviewed and approved by the authors.

---

**Figure S1.**
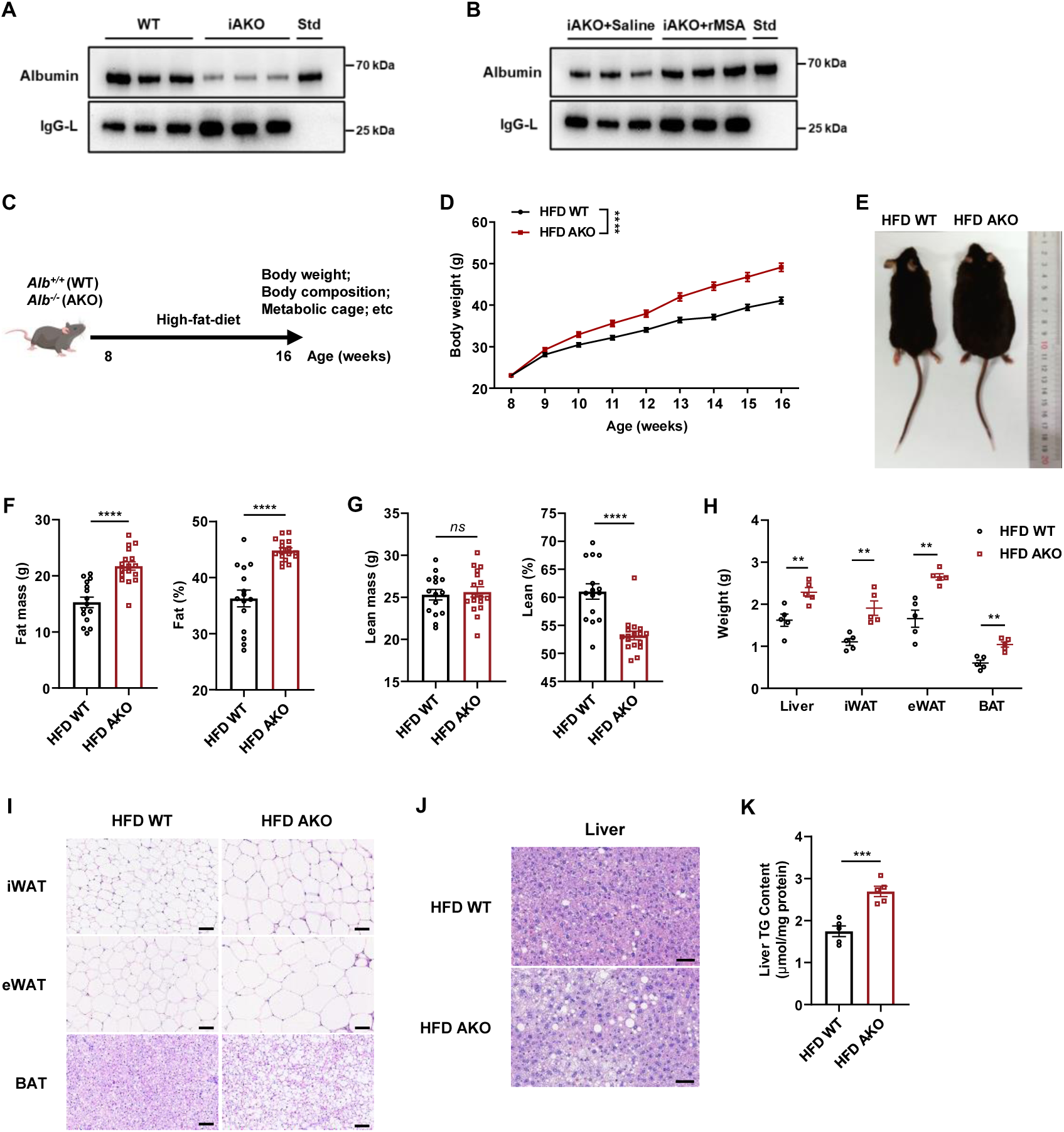
Albumin deficiency exacerbates obesity progression in mice. (A) Serum albumin levels assessed by Western blot (upper). The rightmost lane represents the 30 mg/ml albumin standard. n = 3 biological replicates per group. (B) Serum albumin levels assessed by Western blot (upper). The rightmost lane represents the 30 mg/ml albumin standard. n = 3 biological replicates per group. (C) Schematic illustration of the animal experimental procedure. (D) Body weights of WT and Albumin knockout (AKO) mice on high-fat diet (HFD). n = 22 for WT mice, n = 18 for AKO mice. (E) Representative photographs of WT and AKO mice on HFD. (F) Fat mass (left) and fat percentage (right) of WT and AKO mice on HFD. n =15 for WT mice, n = 17 for AKO mice. (G) Lean mass (left) and lean percentage (right) of WT and AKO mice on HFD. n =15 for WT mice, n = 17 for AKO mice. (H) Weights of the liver and adipose tissues. n = 5 mice per group. (I) Representative H&E staining of dissected adipose tissues. Scale bar, 100 μm. (J) Representative H&E staining of dissected liver tissues. Scale bar, 100 μm. (K) Quantification of liver TG content. n = 5 mice per group. Data are presented as mean ± s.e.m; \**P* < 0.05, \*\**P* < 0.01, \*\*\**P* < 0.001, \*\*\*\**P* < 0.0001 compared to control group. *ns*, not significant; two-tailed Student’s t-tests (F-H, and K), two-way ANOVA followed by Sidak’s multiple comparisons (D).

**Figure S2.**
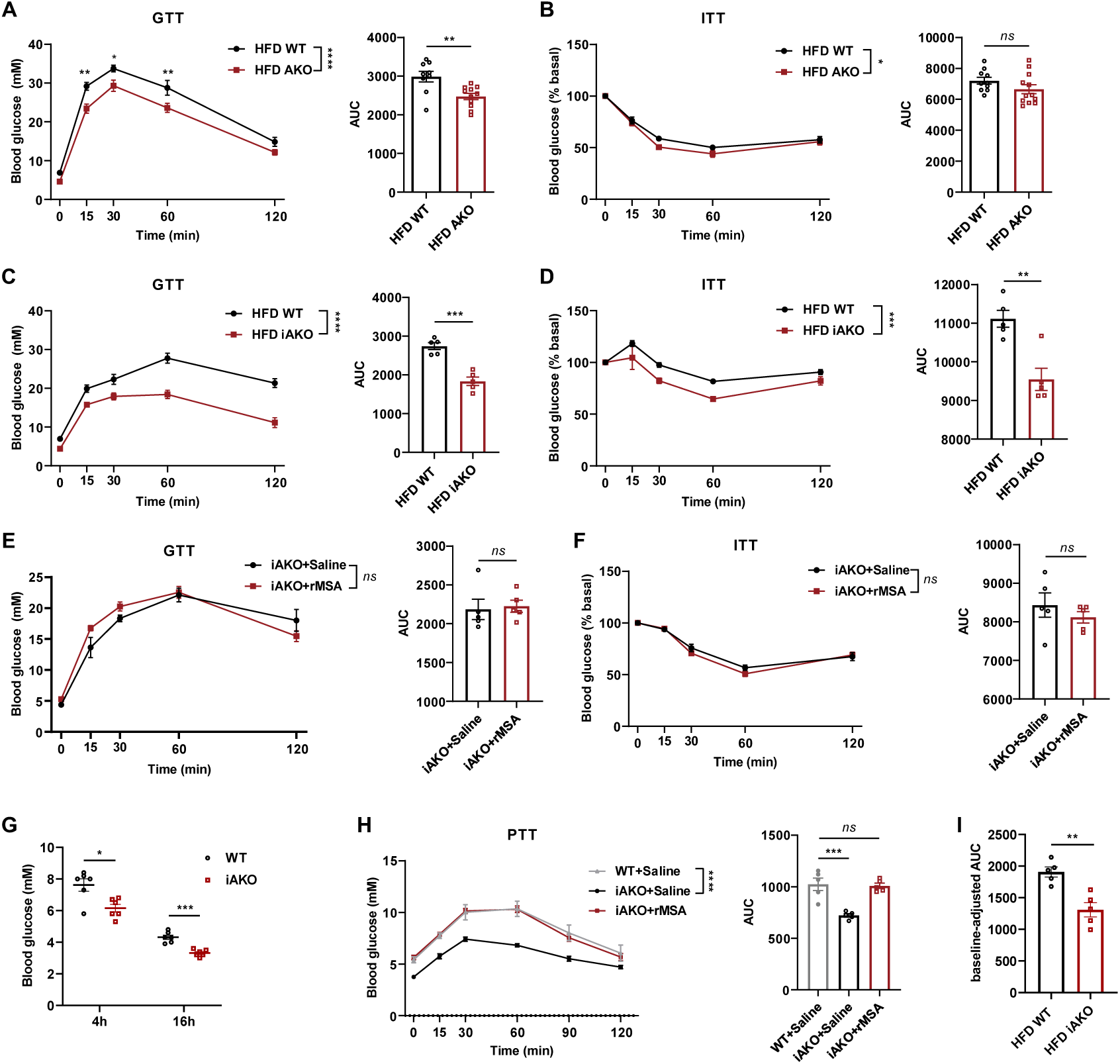
Albumin deficiency alters systemic glucose homeostasis in mice. (A) Glucose tolerance test (GTT) and corresponding area under the curves (AUC) of WT and AKO mice on HFD. n = 9 for WT mice, n = 11 for AKO mice. (B) Insulin tolerance test (ITT) and corresponding AUC of WT and AKO mice on HFD. n = 10 for WT mice, n = 12 for AKO mice. (C) GTT and corresponding AUC of WT and iAKO mice on HFD. n = 5 mice per group. (D) ITT and corresponding AUC of WT and iAKO mice on HFD. n = 5 mice per group. (E) GTT and corresponding AUC of iAKO mice treated with saline or rMSA on HFD. n = 5 mice per group. (F) ITT and corresponding AUC of iAKO mice treated with saline or rMSA on HFD. n = 5 mice per group. (G) Fasting blood glucose levels of WT and AKO mice on normal chow diet. n = 6 mice per group. (H) Pyruvate tolerance test (PTT) and corresponding AUC in WT treated with saline and iAKO mice treated with saline or rMSA on normal chow diet. n = 8 mice per group. (I) Corresponding baseline-adjusted AUC calculated from the GTT data in (C) after subtraction of the 0-min glucose value. n = 5 mice per group. Data are presented as mean ± s.e.m; \**P* < 0.05, \*\**P* < 0.01, \*\*\**P* < 0.001, \*\*\*\**P* < 0.0001 compared to control group. *ns*, not significant; two-tailed Student’s t-tests (A-G), one-way ANOVA (H and I), two-way ANOVA followed by Sidak’s multiple comparisons (A-F, H).

**Figure S3.**
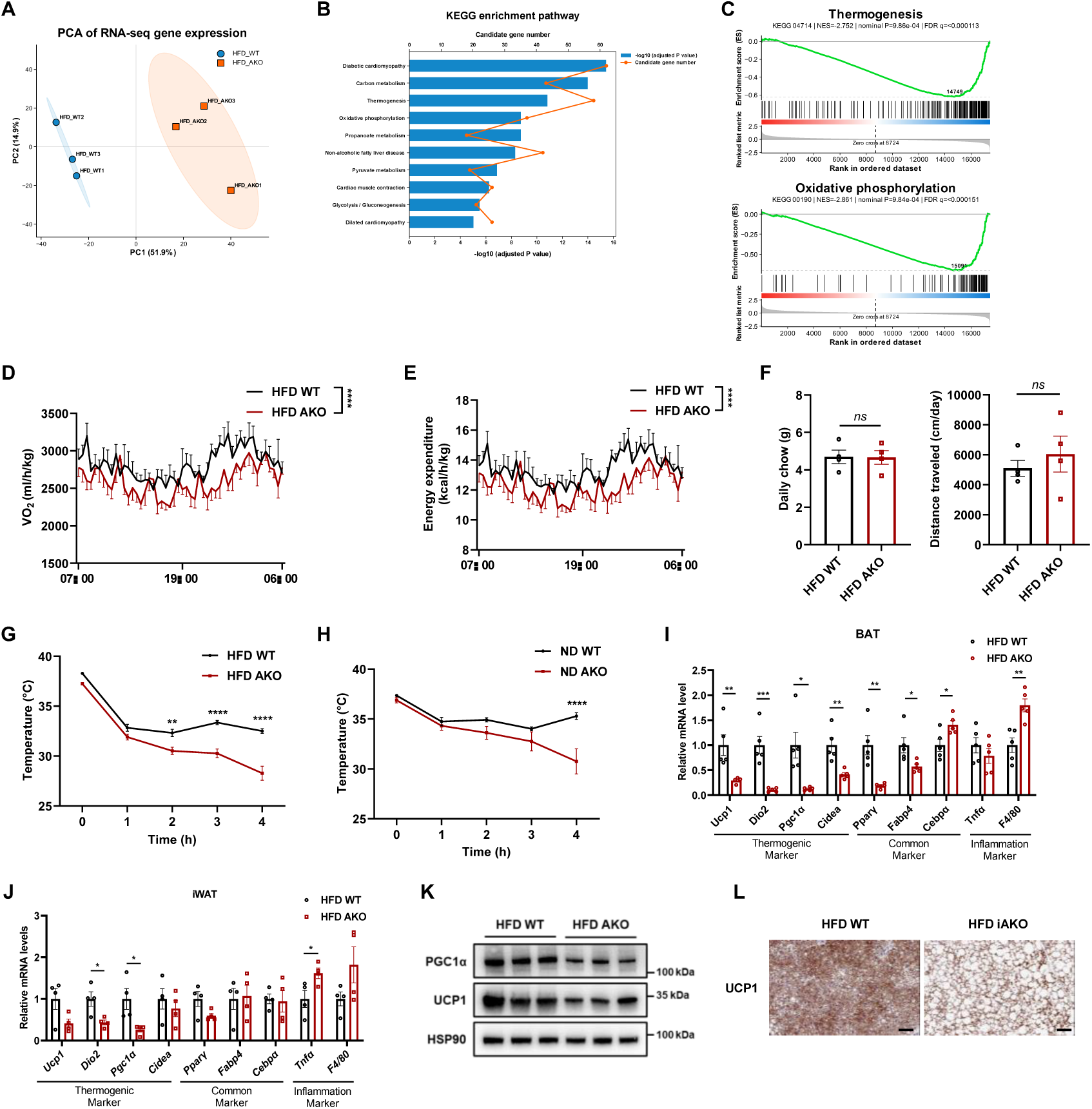
Albumin contributes to the regulation of thermogenesis in adipose tissue. (A) Principal component analysis (PCA) of RNA-sequencing profiles from BAT of HFD-fed WT and AKO mice. (B) KEGG pathway enrichment analysis of genes significantly downregulated in BAT from HFD-fed AKO mice relative to HFD-fed WT mice. Bars indicate −log10(adjusted P value), and orange points and lines indicate the number of genes assigned to each pathway. (C) Gene set enrichment analysis (GSEA) of the KEGG thermogenesis (mmu04714; upper) and oxidative phosphorylation (mmu00190; lower) gene sets in BAT from HFD-fed WT and AKO mice. (D) Whole-body VO_2_ normalized by body weight of WT and AKO mice. n = 4 mice per group. (E) Energy expenditure normalized by body weight of WT and AKO mice. n = 4 mice per group. (F) Daily chow and distance traveled of WT and AKO mice. n = 4 mice per group. (G) Body temperature of WT and AKO mice exposed to cold. n = 10 mice per group. (H) Body temperature of WT and AKO mice on normal diet exposed to cold. n = 5 mice per group. (I) Abundance of thermogenic-related, common adipocyte and inflammation-related mRNAs in BAT. n = 5 mice per group. (J) Abundance of thermogenic-related, common adipocyte and inflammation-related mRNAs in iWAT from HFD WT or AKO mice. n = 4 mice per group. (K) Immunoblots of thermogenesis-related proteins in BAT. n = 3 biological replicates per group. (L) Representative UCP1 immunohistochemical staining in BAT. Scale bar, 100 μm. For RNA sequencing, n = 3 mice per group. Data are presented as mean ± s.e.m; \**P* < 0.05, \*\**P* < 0.01, \*\*\**P* < 0.001, \*\*\*\**P* < 0.0001 compared to control group. *ns*, not significant; two-tailed Student’s t-tests (F, I, and J), two-way ANOVA followed by Sidak’s multiple comparisons (D-E, and G-H).

**Figure S4.**
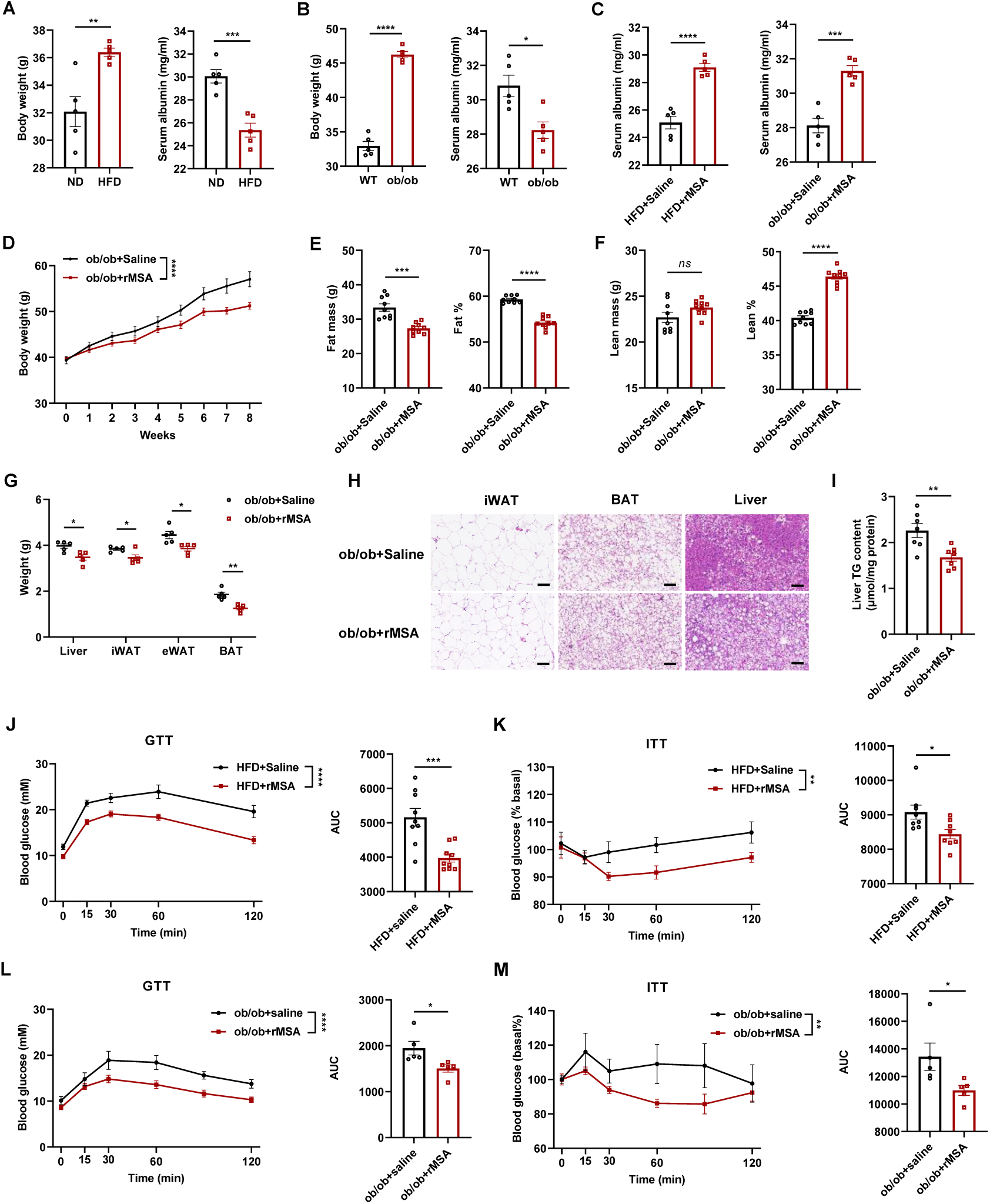
rMSA administration reduces fat mass and improves glucose homeostasis in obese mice. (A) Body weight and serum albumin levels of ND mice and HFD mice. n = 5 mice per group. (B) Body weight and serum albumin levels of WT mice and ob/ob mice. n = 5 mice per group. (C) Serum albumin levels of DIO mice and ob/ob mice after one injection saline or rMSA. n = 5 mice per group. (D) Body weights of ob/ob mice treated with saline or rMSA. n = 9 mice per group. (E) Fat mass (left) and fat percentage (right) of ob/ob mice treated with saline or rMSA. n = 9 mice per group. (F) Lean mass (left) and lean percentage (right) of ob/ob mice treated with saline or rMSA. N = 9 mice per group. (G) Weights of liver and fat tissue. n = 5 mice per group. (H) Representative H&E staining of dissected adipose tissues. Scale bar, 100 μm. (I) Quantification of Liver TG content. n = 7 mice per group. (J) GTT and corresponding AUC in DIO mice treated with saline or rMSA. n = 9 mice per group. (K) ITT and corresponding AUC in DIO mice treated with saline or rMSA. n = 9 mice per group. (L) GTT and corresponding AUC in ob/ob mice treated with saline or rMSA. n = 5 mice per group. (M) ITT and corresponding AUC in ob/ob mice treated with saline or rMSA. n = 5 mice per group. Data are presented as mean ± s.e.m; \**P* < 0.05, \*\**P* < 0.01, \*\*\**P* < 0.001, \*\*\*\**P* < 0.0001 compared to control group. ns, not significant; two-tailed Student’s t-tests (A-C, E-G, and I-M), two-way ANOVA followed by Sidak’s multiple comparisons (D, and J-M).

**Figure S5.**
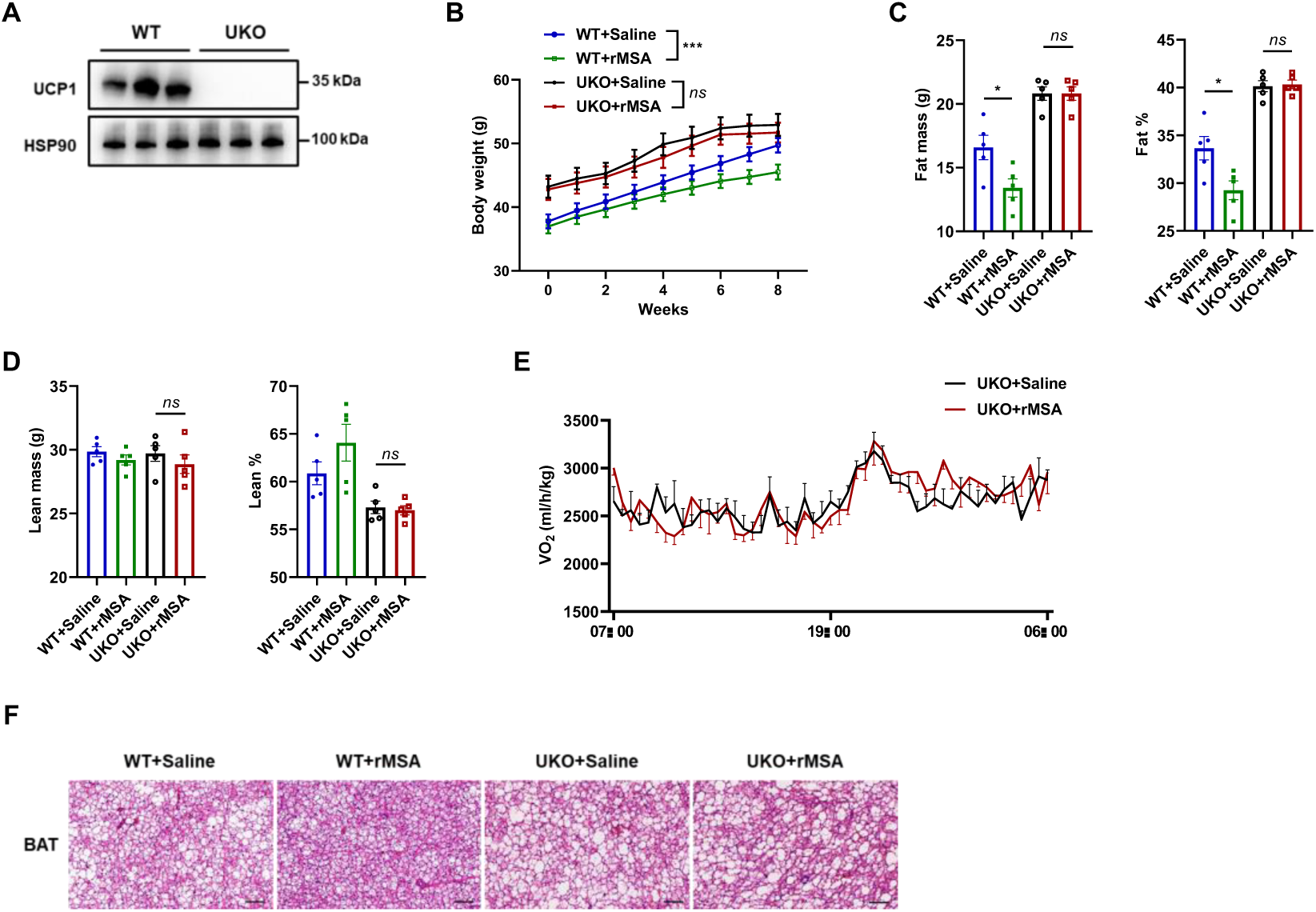
The anti-obesity effects of rMSA are partly dependent on UCP1. (A) Representative immunoblot analysis of UCP1 in BAT from WT and Ucp1-knockout (UKO) mice. HSP90 was used as a loading control. n = 3 biological replicates per group. (B) Body weights of WT and Ucp1-knockout (UKO) mice on HFD treated with saline or rMSA. n = 5 mice per group. For endpoint body weight, the genotype × treatment interaction was not significant (*P* = 0.3094). (C) Fat mass (left) and fat percentage (right) of WT and UKO mice treated with saline or rMSA. n = 5 mice per group. Significant genotype × treatment interactions were detected for fat mass (p = 0.0391) and fat percentage (*P* = 0.0173). (D) Lean mass (left) and lean percentage (right) of WT and UKO mice treated with saline or rMSA. n = 5 mice per group. (E) Whole-body VO_2_ and energy expenditure normalized by body weight of UKO mice. n = 4 mice per group. (F) Representative H&E staining of BAT from WT and UKO mice treated with saline or rMSA. Scale bar, 100 μm. Data are presented as mean ± s.e.m; \**P* < 0.05, \*\**P* < 0.01, \*\*\**P* < 0.001, \*\*\*\**P* < 0.0001 compared to control group. ns, not significant; two-way ANOVA followed by Sidak’s multiple comparisons (B and E). Endpoint body weight, fat mass, fat percentage, lean mass, and lean percentage were analyzed by ordinary two-way ANOVA with genotype (WT versus UKO) and treatment (saline versus rMSA) as factors.

**Figure S6.**
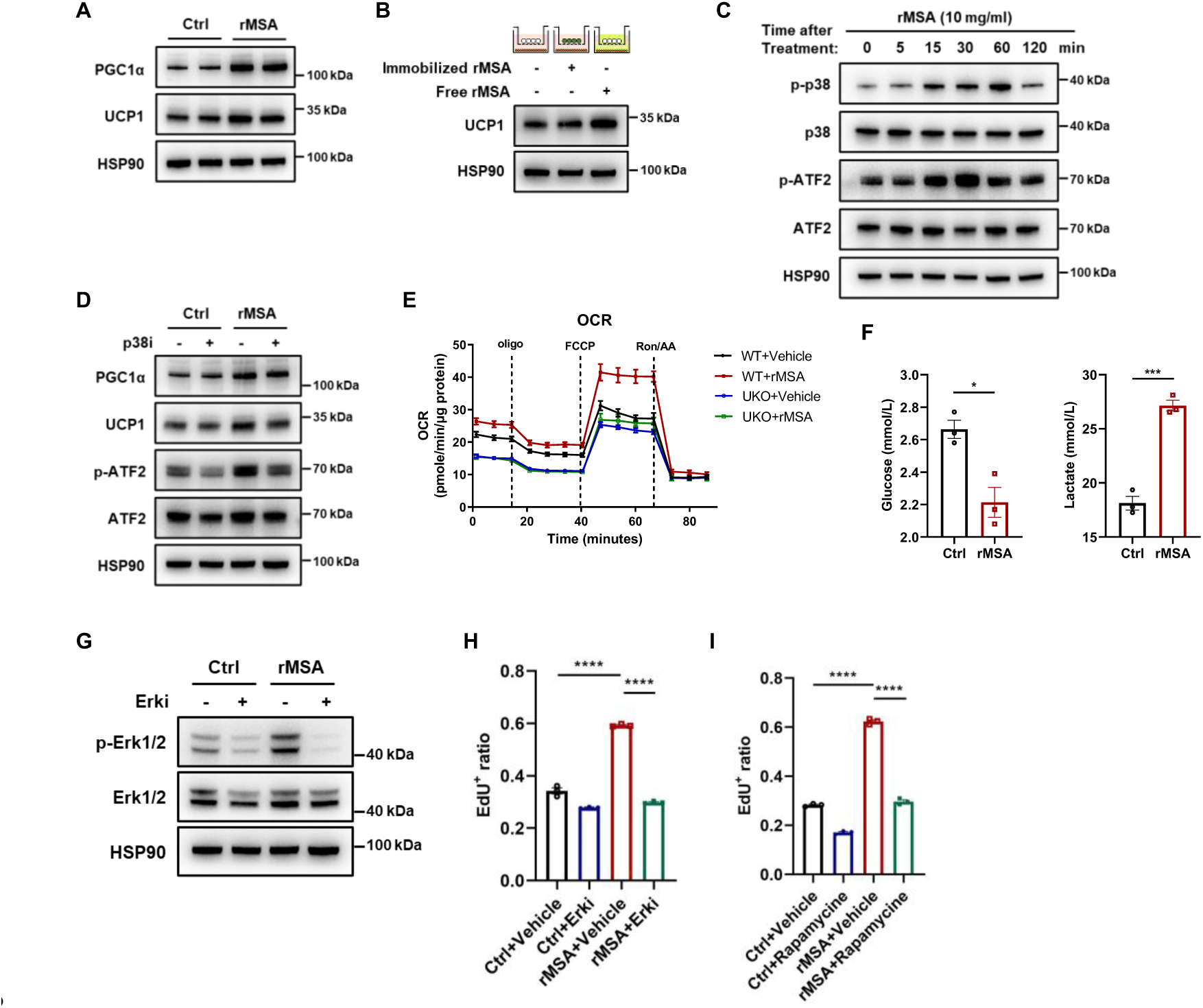
rMSA promotes brown-adipocyte thermogenic activity and preadipocyte proliferation in vitro. (A) Immunoblots (proteins indicated) in control and rMSA-treated mature primary brown adipocytes. (B) Immunoblots (proteins indicated) in agarose beads, immobilized rMSA and free rMSA-treated mature primary brown adipocytes. (C) Immunoblots (proteins indicated) in rMSA treated mature primary brown adipocytes at different time points. (D) Immunoblots (proteins indicated) in mature primary brown adipocytes incubated with or without p38 inhibitor SB203580 (10 μM) for 30 min, followed by rMSA treatment for 12 h. (E) OCR measured by Seahorse mitochondrial stress testing in primary brown adipocytes from WT and UCP1 knockout mice. n = 12 biological replicates per group. (F) Quantification of glucose and lactate concentration from media. The media was collected after incubated with mature brown adipocytes for 48 h. n = 3 biological replicates per group (G) Representative immunoblot analysis of Erk1/2 phosphorylation of brown preadipocytes that were pretreated with or without Erki (SCH772984, 10 μM) and stimulated with or without rMSA. (H) Quantification of primary mouse proliferating cells that were pretreated with or without Erki and stimulated with or without rMSA. n = 3 biological replicates per group. (I) Quantification of primary mouse proliferating cells that were pretreated with or without Rapamycin and stimulated with or without rMSA. n = 3 biological replicates per group. Data are presented as mean ± s.e.m; \**P* < 0.05, \*\**P* < 0.01, \*\*\**P* < 0.001, \*\*\*\**P* < 0.0001 compared to control group. ns, not significant; two-tailed Student’s t-tests (F), two-way ANOVA followed by Sidak’s multiple comparisons (E)

**Figure S7.**
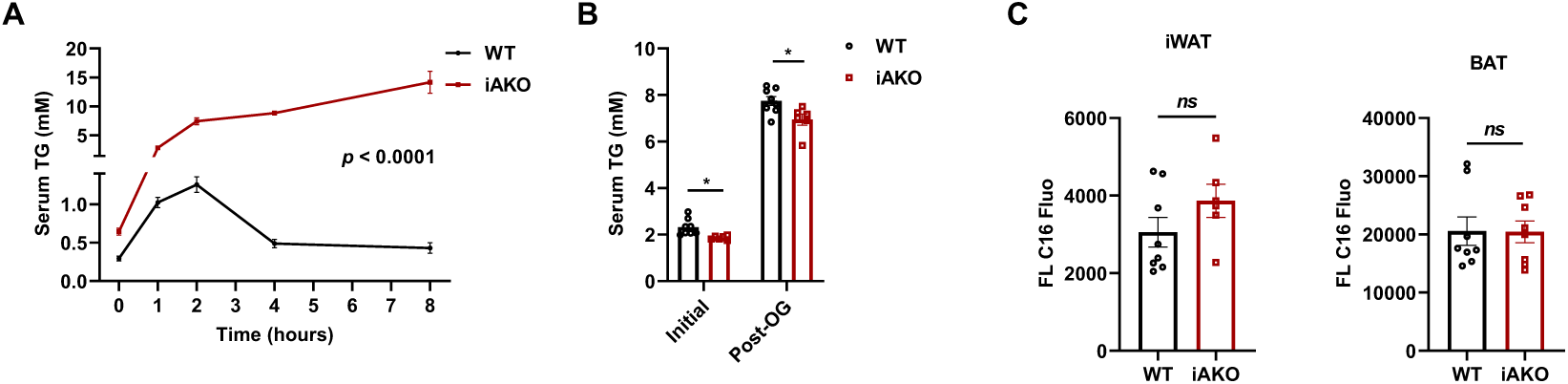
Serum triglyceride dynamics and fatty acid uptake in WT and iAKO mice following oral gavage. (A) Lipid tolerance test on WT and iAKO mice. n = 8 mice for WT, n = 6 for iAKO. (B) Mice were fasted and then given an intraperitoneal injection of poloxamer 407. One hour later, TG levels were measured from the serum and the mice were given an oral olive oil bolus. Two hours later, serum triglyceride levels were measured again. n = 8 mice for WT, n = 6 for iAKO. (C) BODIPY FL C16 in inguinal white adipose tissue (iWAT) and brown adipose tissue (BAT) from WT and iAKO mice. Data are presented as mean ± s.e.m; \**P* < 0.05, \*\**P* < 0.01, \*\*\**P* < 0.001, \*\*\*\**P* < 0.0001 compared to control group. *ns*, not significant; two-tailed Student’s t-tests (B and C), two-way ANOVA followed by Sidak’s multiple comparisons (A).

**Figure S8.**
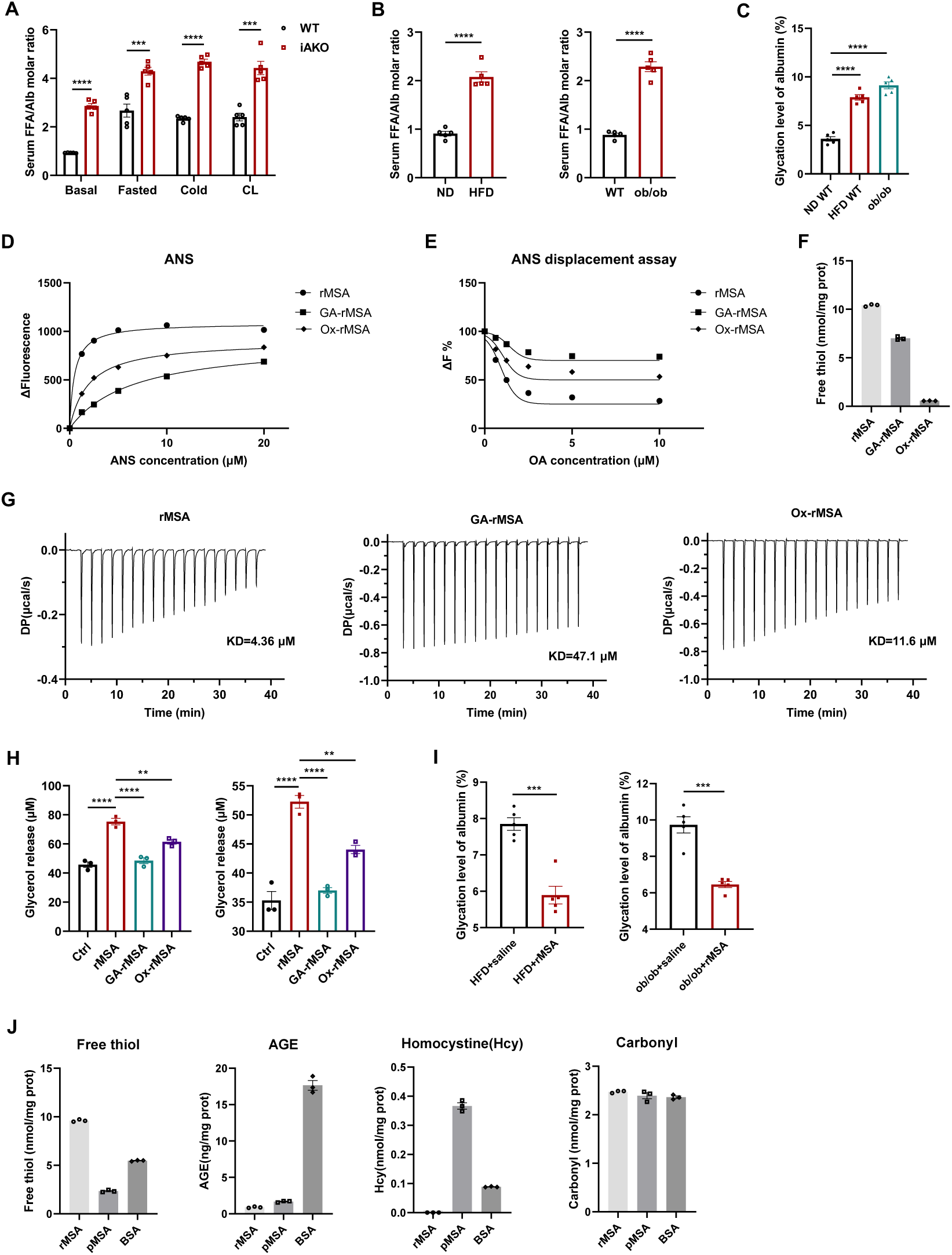
Circulating fatty acid burden relative to albumin and biochemical determinants of albumin fatty-acid-binding capacity. (A) Serum FFA-to-albumin molar ratio in WT and iAKO mice at baseline and following fasting, cold exposure, or CL316243 (CL) stimulation. The molar ratio was calculated from matched serum FFA and albumin concentrations. n = 5 mice per group (B) Serum FFA-to-albumin molar ratio in normal-diet-versus high-fat-diet-fed WT mice (left) and in WT versus ob/ob mice (right). The molar ratio was calculated from matched serum FFA and albumin concentrations. n = 5 mice per group. (C) Glycation levels of circulating albumin from normal-diet-fed WT (ND WT), high-fat-diet-fed WT (HFD WT), and ob/ob mice. n = 5 mice per group. (D) ANS fluorescence assay of recombinant mouse serum albumin (rMSA), glycated rMSA (GA-rMSA), and oxidized rMSA (Ox-rMSA) at the indicated ANS concentrations. (E) ANS displacement assay of rMSA, GA-rMSA, and Ox-rMSA with increasing concentrations of oleic acid (OA). (F) Free thiol contents of rMSA, GA-rMSA, and Ox-rMSA. (G) Representative isothermal titration calorimetry (ITC) thermograms of OA binding to rMSA, GA-rMSA, and Ox-rMSA. The calculated dissociation constants (KD) are indicated. (H) Lipolysis in primary white adipocytes (left) and primary brown adipocytes (right) treated with rMSA, glycated rMSA (GA-rMSA), or oxidized rMSA (Ox-rMSA) for 6 h, as assessed by glycerol release into the culture medium. n = 3 biological replicates per group. (I) Albumin glycation levels in HFD-fed and ob/ob mice treated with saline or rMSA, as indicated. n = 5 mice per group. (J) Free thiol, advanced glycation product (AGE), homocysteine (Hcy), and protein carbonyl contents of rMSA, plasma mouse serum albumin (pMSA), and bovine serum albumin (BSA), as indicated. Data are presented as mean ± s.e.m; \**P* < 0.05, \*\**P* < 0.01, \*\*\**P* < 0.001, \*\*\*\**P* < 0.0001 compared to control group. *ns*, not significant; two-tailed Student’s t-tests (A, B, and I); one-way ANOVA followed by Dunnett’s multiple-comparisons test (C and H).

**Figure S9.**
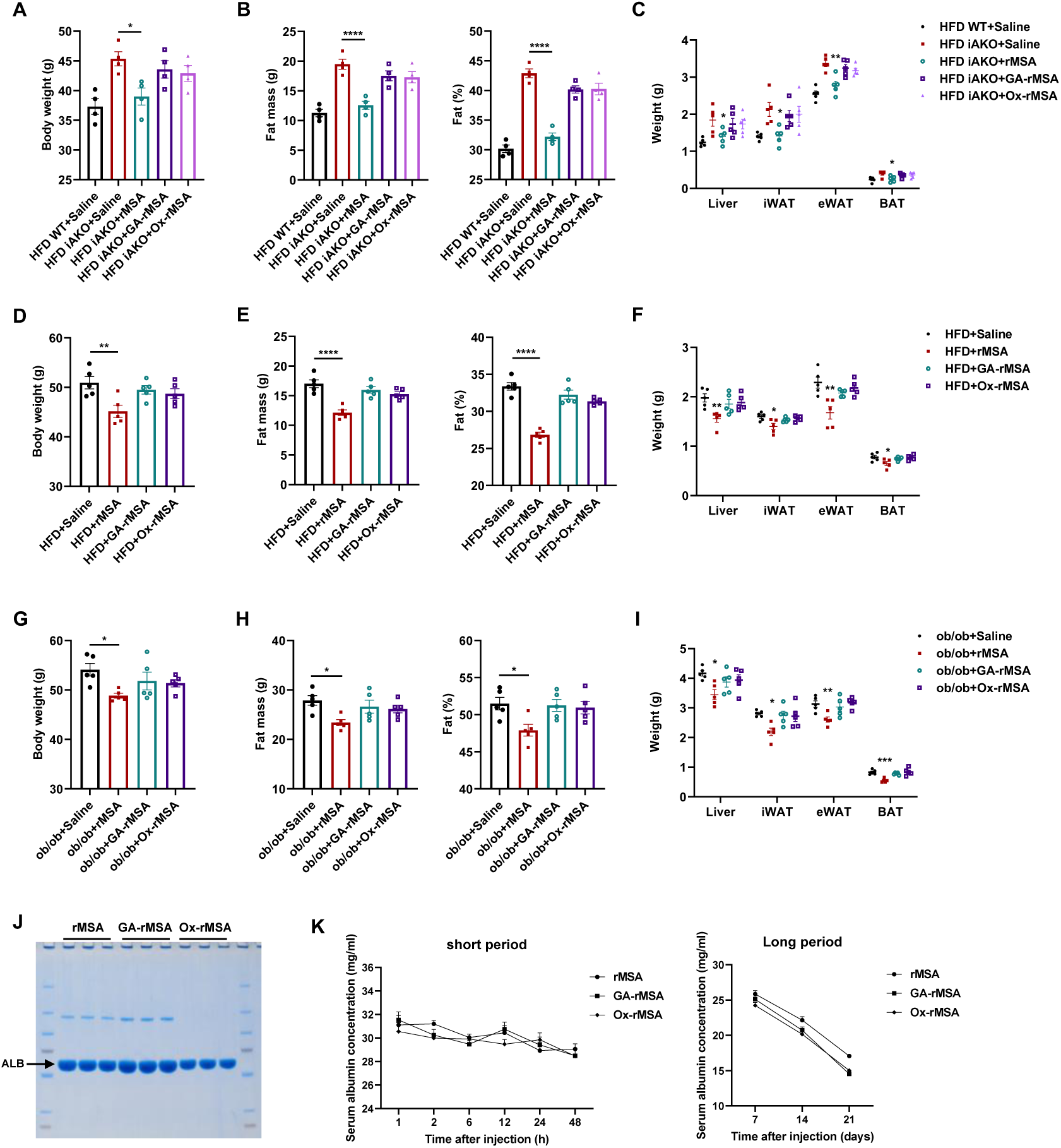
Biochemical modification impairs the anti-obesity activity of rMSA in vivo. (A–C) Body weight (A), fat mass and percentage (B), and liver/adipose-tissue weights (C) in HFD-fed WT mice treated with saline and iAKO mice treated with saline, rMSA, GA-rMSA or Ox-rMSA; n = 5 mice per group. (D–F) Corresponding measurements in DIO mice; n = 5 mice per group. (G–I) Corresponding measurements in ob/ob mice; n = 5 mice per group. (J) Coomassie Brilliant Blue staining of rMSA, GA-rMSA and Ox-rMSA after non-reducing SDS–PAGE. (K) Serum albumin concentrations in AKO mice after a single intravenous injection of rMSA, GA-rMSA or Ox-rMSA (1.5 mg/g), measured by ELISA over 1–48 h (left) and 7–21 d (right); n = 3 mice per group. Data are presented as mean ± s.e.m; \**P* < 0.05, \*\**P* < 0.01, \*\*\**P* < 0.001, \*\*\*\**P* < 0.0001 compared to control group. *ns*, not significant; one-way ANOVA followed by Dunnett’s multiple-comparisons test (A-I). Serum albumin concentrations in (K) were analyzed separately for the short-term and long-term periods using ordinary two-way ANOVA with albumin modification and time as factors. The main effect of albumin modification was not significant during the short-term period (*P* = 0.4082) but was significant during the long-term period (*P* < 0.0001).

