## Supplementary material for "Serum albumin restrains obesity by coordinating brown adipose thermogenic capacity with β3-adrenergic responsiveness": Table S1-S3

**Supplemental Tables**

**Table S1. Antibodies used in this research.**

| Antibodies | SOURCE | IDENTIFIER |
| --- | --- | --- |
| Rabbit monoclonal anti-UCP1 (E9Z2V) | Cell Signaling | Cat#72298; RRID: AB_2936479 |
| Rabbit polyclonal anti-Dio2 | Proteintech | Cat#26513-1-AP; RRID: AB_2880538 |
| Rabbit monoclonal anti- PPAR𝛄 (C26H12) | Cell Signaling | Cat#2435; RRID: AB_ 2166051 |
| Mouse monoclonal anti-HSP90 | ZSGB-Bio | Cat#TA-12; RRID: AB_2106495 |
| Mouse monoclonal anti-β actin | ZSGB-Bio | Cat#TA-09; RRID: AB_2636897 |
| Mouse monoclonal anti-PGC1α (1C1B2) | Proteintech | Cat# 66369-1-Ig; RRID: AB_2828002 |
| Rabbit monoclonal anti-phospho-p38 MAPK (Thr180/Tyr182) (D3F9) | Cell Signaling | Cat#4511; RRID: AB_ 2139682 |
| Rabbit monoclonal anti-p38 MAPK (D13E1) | Cell Signaling | Cat#8690; RRID: AB_10999090 |
| Rabbit monoclonal anti-phospho-Src family (D49G4) | Cell Signaling | Cat#6943; RRID: AB_10013641 |
| Rabbit monoclonal anti-Src (36D10) | Cell Signaling | Cat#2109; RRID: AB_2106059 |
| Rabbit monoclonal anti-ATF2 | Beyotime | Cat#AF1240; RRID: |
| Rabbit monoclonal anti-phospho-ATF2 (Thr71) | Beyotime | Cat#AF1948; RRID: AB_2239535 |
| Rabbit polyclonal anti-phospho-HSL (Ser660) | Cell Signaling | Cat#45804; RRID: AB_2893315 |
| Rabbit monoclonal anti-HSL (D6W5S) | Cell Signaling | Cat#18381; RRID: AB_2798800 |
| Rabbit monoclonal anti-phospho- 4E-BP1 (Thr37/46) (236B4) | Cell Signaling | Cat#2855; RRID: AB_560835 |
| Rabbit monoclonal anti- 4E-BP1 (53H11) | Cell Signaling | Cat#9644; RRID: AB_2097841 |
| Rabbit polyclonal anti-phospho-p70 S6 kinase (Thr389) | Cell Signaling | Cat#9205; RRID: AB_330944 |
| Rabbit polyclonal anti-p70 S6 kinase | Cell Signaling | Cat#9202; RRID: AB_331676 |
| Rabbit monoclonal anti-caveolin-1 (D46G3) | Cell Signaling | Cat#3267; RRID: AB_2275453 |
| Rabbit monoclonal anti-phospho-PKA substrate (RRXS*/T*) (100G7E) | Cell Signaling | Cat#9624; RRID: AB_331817 |
| Rabbit monoclonal anti-phospho-Akt (Ser473) (D9E) | Cell Signaling | Cat#4060; RRID: AB_2315049 |
| Rabbit polyclonal anti-Akt | Cell Signaling | Cat#9272; RRID: AB_329827 |
| Rabbit monoclonal anti-PKM1/2 (C103A3) | Cell Signaling | Cat#3190; RRID: AB_2163695 |
| Mouse monoclonal anti-PFKL (1H2G2) | Proteintech | Cat#68385-1-Ig; RRID: AB_3085103 |
| Rabbit polyclonal anti-HK2 | Proteintech | Cat#22029-1-AP; RRID: AB_11182717 |
| Goat anti-mouse IgG, HRP-linked | Cell Signaling | Cat#7076; RRID: AB_330924 |
| Goat anti-rabbit IgG, HRP-linked | Cell Signaling | Cat#7074; RRID: AB_2099233 |
| Albumin | Proteintech | Cat#16475-1-AP; RRID: AB_2242567 |

**Table S2. Primer sequences for qRT-PCR analysis.**

| Gene | Forward (5’-3’) | Reverse (5’-3’) |
| --- | --- | --- |
| *Ucp1* | CAAAAACAGAAGGATTGCCGAAA | TCTTGGACTGAGTCGTAGAGG |
| *Dio2* | GAAGGCTGCCGAATGTCAAC | GCGTGAGCAGTCTTGTTTGG |
| *Pgc1α* | TATGGAGTGACATAGAGTGTGCT | CCACTTCAATCCACCCAGAAAG |
| *Pparγ* | TCGCTGATGCACTGCCTATG | GAGAGGTCCACAGAGCTGATT |
| *Cidea* | ATCACAACTGGCCTGGTTACG | TACTACCCGGTGTCCATTTCT |
| *Fabp4* | GATGCCTTTGTGGGAACCT | CTGTCGTCTGCGGTGATTT |
| *Tnfα* | TGGGCCTCTCATGCACCACC | GAGGCAACCTGACCACTCTCCCT |
| *F4/80* | CTTTGGCTATGGGCTTCCAGTC | GCAAGGAGGACAGAGTTTATCGTG |
| *Cebpα* | CAAGAACAGCAACGAGTACCG | GTCACTCGTCAACTCCAGCAC |
| *Actb* | GGCTGTATTCCCCTCCATCG | CCAGTTGGTAACAATGCCATGT |

**Table S3. The targeted sequences of siRNAs.**

| Target | Sense (5’-3’) | Antisense (5’-3’) |
| --- | --- | --- |
| Mouse Cav1 #1 | CUGCGAUCCACUCUUUGAATT | AAUUCAAAGAGUGGAUCGCAG |
| Mouse Cav1 #2 | CGCUUGUUGUCUACGAUCUTT | AAAGAUCGUAGACAACAAGCG |
| Mouse CD36 #1 | GUGGGAAACGAUGUCAUUAAA | CACCCUUUGCUACAGUAAUUU |
| Mouse CD36 #2 | ACCAUUGUUUCUGCACUGAAA | UGGUAACAAAGACGUGACUUU |
| Control | UUCUCCGAACGUGUCACGUTT | AAACGUGACACGUUCGGAGAA |
